# Individualized event structure drives individual differences in whole-brain functional connectivity

**DOI:** 10.1101/2021.03.12.435168

**Authors:** Richard F. Betzel, Sarah A. Cutts, Sarah Greenwell, Joshua Faskowitz, Olaf Sporns

## Abstract

Resting-state functional connectivity is typically modeled as the correlation structure of whole-brain regional activity. It is studied widely, both to gain insight into the brain’s intrinsic organization but also to develop markers sensitive to changes in an individual’s cognitive, clinical, and developmental state. Despite this, the origins and drivers of functional connectivity, especially at the level of densely sampled individuals, remain elusive. Here, we leverage novel methodology to decompose functional connectivity into its precise framewise contributions. Using two dense sampling datasets, we investigate the origins of individualized functional connectivity, focusing specifically on the role of brain network “events” – short-lived and peaked patterns of high-amplitude cofluctuations. Here, we develop a statistical test to identify events in empirical recordings. We show that the patterns of cofluctuation expressed during events are repeated across multiple scans of the same individual and represent idiosyncratic variants of template patterns that are expressed at the group level. Lastly, we propose a simple model of functional connectivity based on event cofluctuations, demonstrating that group-averaged cofluctuations are suboptimal for explaining participant-specific connectivity. Our work complements recent studies implicating brief instants of high-amplitude cofluctuations as the primary drivers of static, whole-brain functional connectivity. Our work also extends those studies, demonstrating that cofluctuations during events are individualized, positing a dynamic basis for functional connectivity.

## INTRODUCTION

Functional connectivity (FC) measures the temporal correlation of regional BOLD activity, often in the absence of explicit task instructions, i.e. in the “resting state” [1, 2]. Although usually estimated over an extended period of time and using all available data, a growing number of studies have shown that FC can be well approximated using relatively few observations, suggesting that FC may be driven by a temporally sparse process [3–7].

In parallel, a growing body of work has demonstrated that, like fingerprints, FC is unique to each individual and expresses features that reliably distinguish one brain from another [8–12]. These observations hold tremendous translational promise, and open up the possibility of designing personalized interventions [13] and developing increasingly potent connectivity-based biomarkers for cognition, development, and disease [14–16].

However, there remains a key open question: how does FC become individualized in the first place? One possibility is that, like FC itself, personalized information is encoded through time-varying connectivity patterns and distributed dynamically and sparsely throughout a scan session. Indeed, recent findings broadly support this hypothesis [17, 18]. In [19], for instance, we demonstrated that using a small subset of frames classified as “events - brief and infrequent periods of high-amplitude cofluctuation – we could produce accurate reconstructions of FC while simultaneously rendering participants identifiable, amplifying their connectional fingerprints. In contrast, low-amplitude frames yielded poorer estimates of FC and contained little personalized information.

Although these observations support the hypothesis that personalized information is expressed selectively during high-amplitude frames, they also raise additional theoretical questions (Fig. 1). For instance, do cofluctuation patterns during events repeat from one scan to another (Fig. 1b)? If so, do they reflect a single repeating pattern or a repertoire of different patterns? Are these patterns shared across individuals but expressed in different proportions, thereby giving rise to individualized FC (Fig. 1c)? Or does the individualization of FC arise from equally idiosyncratic patterns of high-amplitude cofluctuations (Fig. 1d)? Addressing these questions is critical for linking patterns of brain connectivity with individual differences in behavior [20], and would help clarify the role of brain dynamics in shaping the individualization of FC [21], complementing other approaches that have focused on the collective influence of cortical expansion rates, post-natal experience, and genetics [22].

**FIG. 1.**
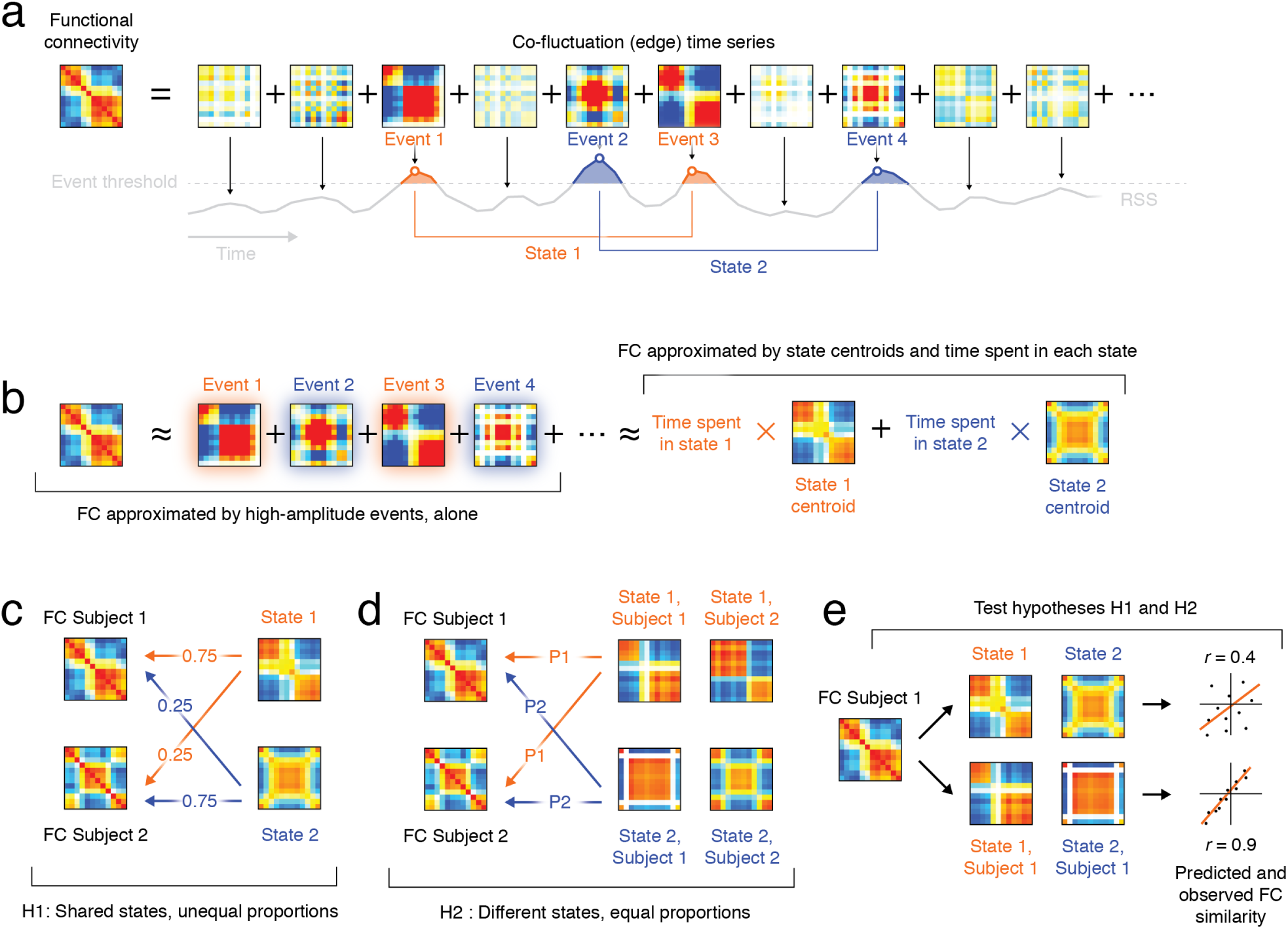
Competing hypotheses for how co-fluctuations contribute to the individualization of FC. (*a*) Edge time series decompose FC into its framewise contributions. (*b*) FC can be well-approximated from co-fluctuations expressed during high-amplitude “events”. Recurrences of event co-fluctuation patterns can be grouped into clusters or “states”. FC can then be approximated from state centroids and the relative frequency with which each state is visited. Why does FC differ between individuals? (*c*) One hypothesis (H1) is that the same states are shared across individuals and inter-individual differences are driven by differences in the frequency with which those shared states are visited. (*d*) Another hypothesis (H2) is that the states, themselves, are subject-specific. In this case, inter-individual differences in FC are driven by differences in the state centroids across subjects. (*e*) To adjudicate between these hypotheses, we can try to approximate FC matrices with centroids estimated from that same subject’s data (different scans) or from group-averaged data.

Here, we address these questions directly. Our approach leverages a recently-proposed method for decomposing FC into its framewise contributions, detecting events, and assessing the impact of events on time-averaged FC [19, 23–30]. We apply this framework to two independently acquired datasets: the Midnight Scan Club [11, 12] and the MyConnectome project [10, 31]. In agreement with our previous studies, we show that FC is accurately reconstructed from event data alone. Next, we focus on the properties of individual events, revealing that they repeat within and between scans of the same individual. We also show that event cofluctuations can be clustered across participants, revealing broad archetypes that are subtly yet systematically modified at the level of individuals. Finally, we construct a simple model of FC, demonstrating that FC can be predicted with a high level of accuracy using individualized event data, exclusively.

## RESULTS

### Edge time series as a mathematical precise link between brain dynamics and FC

In this paper, we analyze data from eight participants in the Midnight Scan Club, each scanned ten times (participants MSC08 and MSC09 were dropped due to data quality issues). We analyzed two versions of these data; one in which participants’ brains were parcellated into *N* = 333 group-level parcels [32] and another in which parcels were defined on an individual basis, resulting in a different set parcels for each participant (*N* = 612 ± 28) [33]. The primary analyses were carried out using the group-level parcels. We also analyzed data from the MyConnectome project, a study in which a single individual was scanned > 100 times [10, 31].

For each dataset, we transformed regional fMRI BOLD time series into cofluctuation or edge time series (ETS). Briefly, ETS are calculated as the element-wise product between pairs of z-scored regional (nodal) time series (Fig. 2a; see **Materials and Methods** for details). This operation results in a new time series – one for every node pair (edge) – whose elements index the direction and magnitude of instantaneous cofluctuations between the corresponding pair of brain regions. For instance, if the activity of region *i* and *j* deflect above (or below) their time-averaged means at the same instant, the value of the edge time series will be positive. On the other hand, if they deflect in opposite directions, then the edge time series returns a negative value. If one deflects and the other does not, then the value will be close to zero. The temporal mean of an edge time series is equal to the Pearson sample correlation coefficient, and therefore ETS is an exact decomposition of FC into its framewise contributions.

**FIG. 2.**
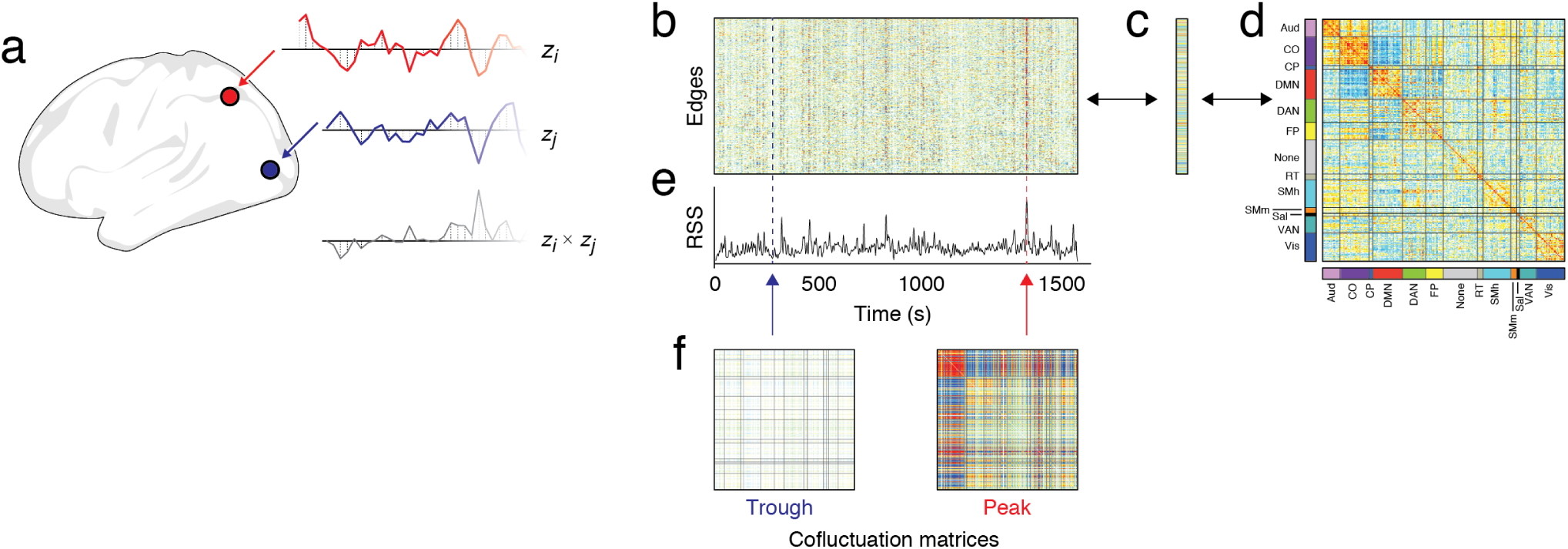
Edge time series. (*a*) An edge time series is constructed for pairs of brain regions, *i* and *j*, by computing the elementwise product of their z-scored activities, *z*_*i*_ and *z*_*j*_, respectively. The result is a new time series, *z*_*ij*_, which indexes the framewise cofluctuations between *i* and *j*. (*b*) This procedure can be repeated for all pairs of regions, generating a matrix of edge time series. At each instant in time, a “slice” through this matrix yields a region-by-region cofluctuation matrix that can be modeled as a network. (*c*) At every moment in time we can calculate the root sum of squares (RSS) over all edge time series. The RSS time series is bursty, such that it takes on low values most of the time, but is punctuated by short, intermittent, high-amplitude bursts. (*d*) The temporal average over all edge time series yields a vector that corresponds to the upper triangle elements of a correlation matrix, i.e. functional connectivity (*e*). In this way, edge time series offer a means of tracking moment-to-moment fluctuations in network topology and links them to functional connectivity through an exact decomposition. In *f* we show examples of cofluctuations during a trough and peak (when RSS is small *versus* large).

If we calculate edge time series for all pairs of regions, we obtain an edge×time matrix (Fig. 2b) whose temporal average yields a vector (Fig. 2c) that, when reshaped into the upper triangle elements of a node×node matrix, is exactly the FC matrix (Fig. 2d). The edge×time matrix can also be “sliced” temporally and the corresponding vector once again reshaped into the upper triangle elements of a node×node matrix, yielding an instantaneous estimate of whole-brain cofluctuations. These matrices vary in terms of their mean cofluctuations, which we summarize with a root sum of squares measure (Fig. 2e).

In our previous study, we showed that RSS values followed a heavy-tailed distribution, such that a small number of frames exhibited exceptionally high-amplitude RSS [19]. We also demonstrated that FC reconstructed using only these high-amplitude frames accurately recapitulated time-averaged FC, suggesting that FC weights are not driven equally by all frames, but by a select set of frames. We also demonstrated that these high-amplitude frames were underpinned by a principal mode of brain activity, emphasizing oppositional activation of default mode and control networks with sensorimotor and attentional networks. In this paper, however, high-amplitude frames were selected heuristically as the top *P*% by RSS value and, beyond the first mode of activity, we did not investigate other activity patterns that occur during events.

### A statistical test for high-amplitude cofluctuation events

In previous work, we identified putative cofluctuation events as the top *P*% frames in terms of root sum squared (RSS) amplitude of cofluctuation weights. Although this heuristic is pragmatic – it is easy to implement and interpret – it has some unwanted characteristics. Notably, the parameter *P*% lacks statistical justification and, due to slow temporal fluctuations and serial correlations in the fMRI BOLD signal, can result in event samples that disproportionately represent only a small number of RSS peaks. Here, we present a simple statistical test to identify events that addresses both of these issues.

In essence, we identify high-amplitude frames by comparing the RSS time series estimated using real data with an ensemble of RSS time series generated under a null model. Here, as a null model we apply the circular shift operator independently, randomly, and bidirectionally to each region’s time series, which exactly preserves its mean and variance (and its autocorrelation approximately). We then transform the shifted data into edge time series and estimate their RSS. This step is repeated 100 times yielding 100 sets of surrogate RSS time series, against which we compare the observed RSS data and identify sequences of frames whose RSS exceeds the null distribution (non-parametric permutation test at each frame; accepted false discovery rate fixed at *q* = 0.05; Figure. 3a). This entire procedure is repeated for every participant and every scan.

**FIG. 3.**
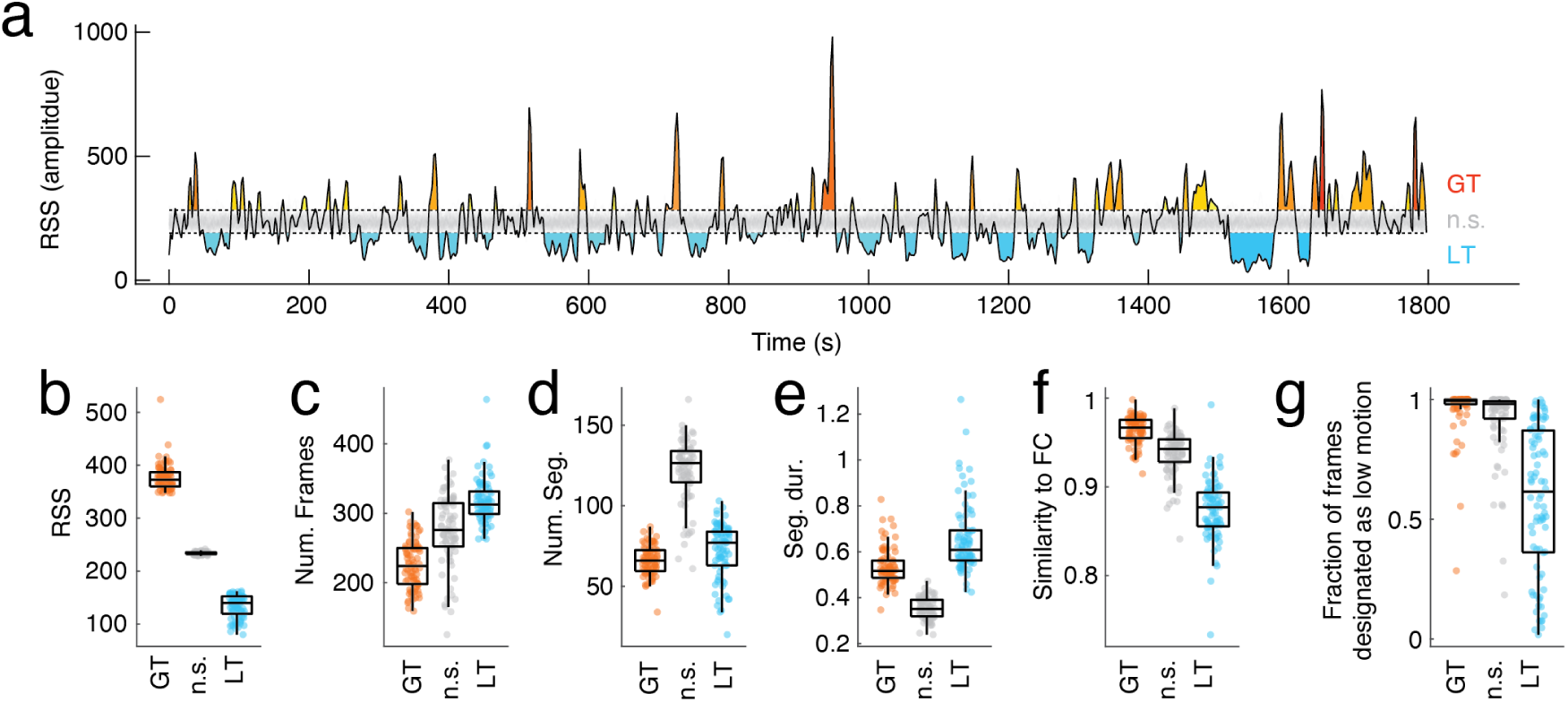
A statistical test for network-wide events. (*a*) We generated edge time series and computed the RSS time series. We compared this time series to a null RSS time series estimated from edge time series that had been generated after circularly shifting the original fMRI BOLD time series. At each point in time, we calculated the probability that the observed RSS value exceeded the null distribution, controlled for multiple comparisons, and identified sequences of frames that exceeded the null distribution. This allowed us to categorize time points into three classes: those whose RSS was greater than null (GT), those whose RSS was significantly less than the null (LT), and those that were in between (n.s.). For subsequent analysis, we extracted a single representative cofluctuation pattern for every contiguous sequence of frames that was greater/less than the null distribution. This pattern corresponded to the frame with the maximum/minimum RSS. In panels *b*-*g*, we separate the frames of each scan into these three classes and compare their features to one another. Each point represents the mean value over all frames assigned to a given class. The features that are compared are: *b* mean RSS, *c*, the number of frames assigned to a given class, *d* the number of contiguous sequences of each class, *e* the mean duration of sequences (*log*_10_ transformed), *f* the similarity of time-average FC with FC reconstructed using only frames assigned to each class, and *g* the fraction of frames censored for high levels of in-scanner motion.

This procedure allows us to segment the time series into three categories: contiguous frames whose RSS is greater than expected, less than expected, or consistent with that of the null distribution. Rather than consider all frames, we select representative frames from each block for subsequent analysis. For segments whose RSS is greater than that of the null distribution or not significant, we extract peak cofluctuation pattern corresponding to the maximum RSS frame; for segments whose RSS is less than that of the null, we extract the pattern corresponding to the minimum RSS frame (trough).

To demonstrate that these categories of frames capture distinct features of cofluctuations, we compare them along several different dimensions (ANOVA; for all comparisons *p <* 10^*−*15^). First, we show that, as expected, high-amplitude frames express greater RSS values than low-amplitude and non-significant frames (Fig. 3b). On the other hand, the number of high-amplitude frames in a scan is smaller than the number of low-amplitude frames (Fig. 3c). Additionally, high-amplitude frames form fewer contiguous segments than low-amplitude frames (Fig. 3d), and, when they do, those segments tend to be of shorter duration then contiguous segments of low-amplitude frames (Fig. 3e). Consistent with our previous study, reconstructing FC using only high-amplitude frames results in a pattern of FC strongly correlated with the FC estimated using all frames, and greater in magnitude than that of the non-significant and low-amplitude frames (Fig. 3f). We note, however, that here the gap in correlation between the high- and low-amplitude is narrower than in previous studies [19]. This is due to differences in the total number of frames used to reconstruct FC and how those frames were selected (see Fig. S1 for more details). Lastly, we find that high-amplitude frames are almost never among those censored for excessive in-scanner motion (Fig. 3g); low-amplitude frames, on the other hand, were more likely to be associated with censored frames, but were also more variable, a result that can be attributed, at least in part, to stable inter-individual differences in motion (Fig. S2). This observa-tion is consistent with our previous study, in which we reported a weak but consistent negative correlation between RSS and framewise displacement [19]. These findings, in general, replicate using individualized parcels for participants in the Midnight Scan Club (Fig. S3) and with MyConnectome data (Fig. S4).

Taken together, these results suggest that the proposed statistical test segments frames into classes with distinct features. This test addresses two concerns associated with previous estimates of high-amplitude “event” frames. First, it defines high-amplitude frames according to a statistical criterion, rather than heuristically. Second, by extracting representative frames from each contiguous segment, we obtain a more heterogeneous sample of high-amplitude frames and avoid selecting multiple frames from around a single peak.

### Peak cofluctuation patterns are repeated across scan sessions – troughs are not

In the previous section we presented a simple method for estimating statistically significant cofluctuation events and demonstrated that the peak and trough frames of these segments exhibit distinct spatiotemporal properties. Here, we investigate representative cofluctuation patterns extracted from blocks of high- and low-amplitude frames. We first compare the similarity of these cofluctuation patterns, first within-individuals and later between. Then, we present evidence that cofluctuation patterns expressed during the peaks of high-amplitude events recur across scans of the same individual and that these patterns exhibit subject-specificity.

First, we applied the statistical test to MSC scans (excluding MSC08 and MSC09 due to data quality issues; see **Materials and Methods** for more details). We found that each scan included 65.9 ± 9.2 and 72.2 ± 17.1 high- and low-amplitude segments, respectively (paired sample t-test; *p* = 0.0017; *t*(79) = 3.24). After additional quality control in which we excluded segments that included any motion-censored frames, the number of segments whose RSS was significantly greater than the null changed little (61.26 ± 14.7). However, the number of segments with lower-than-expected RSS was reduced dramatically (52.5 ±23.9), reflecting the fact that those frames often coincide with periods of excessive in-scanner motion.

Next, we calculated the spatial similarity of motion-free cofluctuation patterns extracted during RSS peaks and troughs (periods when the RSS was significantly greater or less than the null model; labeled G.T. and L.T. in Fig. 3). We performed this analysis separately for each subject, resulting in eight similarity (correlation) matrices. We grouped these values based on whether similarity was measured between two peaks, two troughs, or a peak and trough co-fluctuation pattern. We found peak-peak similarity was significantly greater than trough-trough and peak-trough (t-test; *p <* 10^*−*15^) (Fig. 4a,b). We see an identical effect in the MyConnectome data (Fig. S5a-c) and when parcels are individualized (Fig. S5d). These observations suggest that high-amplitude events encode subject-specific patterns of cofluctuations.

**FIG. 4.**
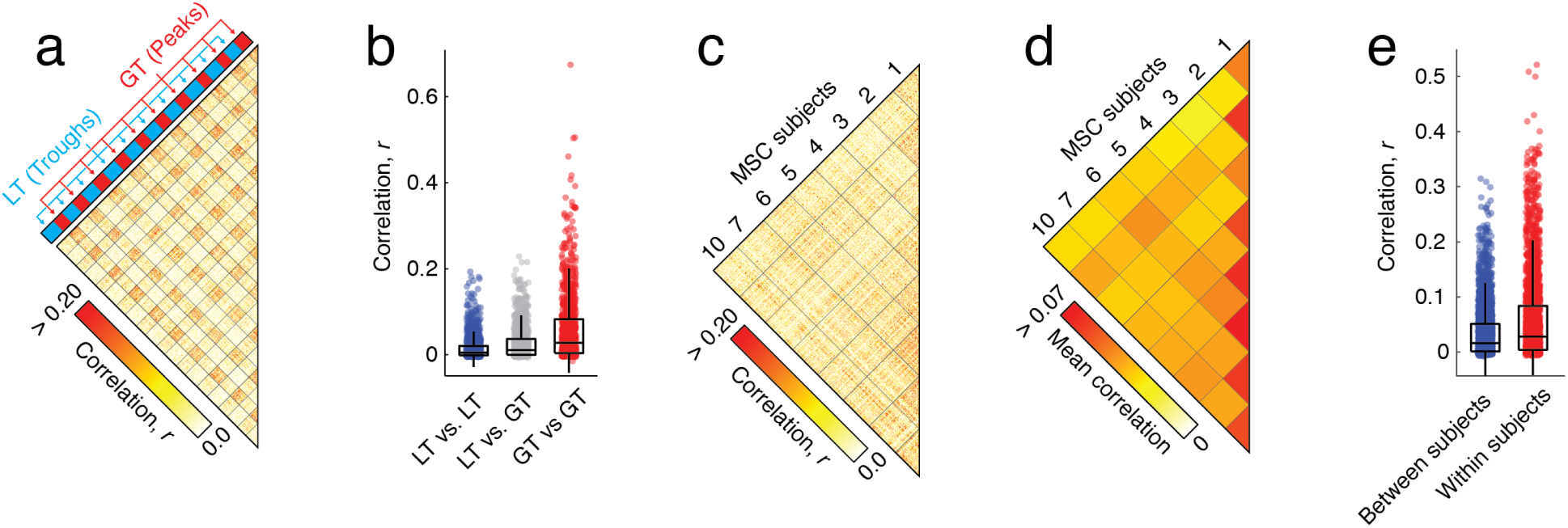
Intra-/Inter-individual similarity of cofluctuation patterns. (*a*) For each individual separately, we aggregated all low-motion cofluctuation patterns during peaks and troughs. We then computed the similarity between cofluctuation patterns (data from participant MSC06 is shown here as an example). (*b*) Boxplot showing similarity values broken down by trough *versus* trough, peak *versus* trough, and peak *versus* peak. Because only cofluctuation at peaks exhibited similarity across scans, we focused on these patterns only, discarding cofluctuation that occurs during troughs and focusing on comparisons of participants to one another. We found that cofluctuation at peaks tended to be more similar within participants than between. We show the raw similarity matrix in *c* and the averaged values in *d*. (*e*) Boxplot of similarity values broken down according to whether they fell within or between participants.

Based on these observations, along with the fact that high-amplitude events are less likely to be impacted by motion, we calculated the spatial similarity of peak cofluctuation patterns for all pairs of detected events, for all scans, and for all subjects (Fig. 4c). We then compared these similarity values based on whether they came from the same or different participants (Fig. 4d). We found that within-individual similarity exceeded between-individual similarity (non-parametric permutation test, *p <* 10^*−*15^; Fig. 4f).

Collectively, these results are in line with our previous study and suggest that low-amplitude cofluctuations contains little participant-specific information [19]. Rather, our findings support the hypothesis that high-amplitude cofluctuations contribute significantly more information about an individual than low-amplitude cofluctuations.

### High-amplitude events can be divided into distinct communities based on their cofluctuation patterns

In the previous section we found that co-fluctuation patterns expressed during peaks of high-amplitude event segments are not related to motion and that they are repeatable across scans. This is in contrast to cofluctuation patterns expressed during low-amplitude segments, which tend to coincide with excessive in-scanner motion and are dissimilar across scans and individuals, even within the same scan session. These observations motivated us to focus on high-amplitude events in yet greater detail. In this section, we test whether the cofluctuation patterns expressed during high-amplitude events are entirely subject-specific and not shared across individuals or whether they belong to a general archetype that is fine-tuned to single participants.

To test whether this is the case, we aggregated across participants all cofluctuation patterns that occurred during event peaks, calculated the similarity matrix of those patterns, and clustered this matrix using a variant of multi-resolution consensus clustering (modularity maximization with a uniform null model [34, 35]; see **Materials and Methods** for details). The results of this analysis yielded two large communities (clusters) along with many very small communities. We found that every participant was represented in the two large communities (labeled 1 and 2 in Fig. 5a) and that instances of those communities appeared in every scan session, accounting for 54.1% and 19.0% of all event peaks, respectively (see Fig. S6). Every participant was also represented in the two next-largest communities, although they appeared infrequently across scan sessions and collectively accounted for only 8.6% of all event peaks. Accordingly, we aggregated the smaller communities to form a third larger community (labeled 3 in Fig. 5a). Most subsequent analyses will focus on communities 1 and 2 unless otherwise noted. For completeness, we analyze community 3 in greater detail in the **Supplementary Material** (see Fig. S7 and Fig. S8). Note that we also repeated this clustering analysis for each participant individually and found that subject-level partitions of events were highly similar to partitions estimated with the group-aggregated data (mean±standard deviation adjusted Rand index across subjects of 0.81 ± 0.10; *p <* 10^*−*4^, permutation test in which each subject’s community labels were randomly shuffled).

**FIG. 5.**
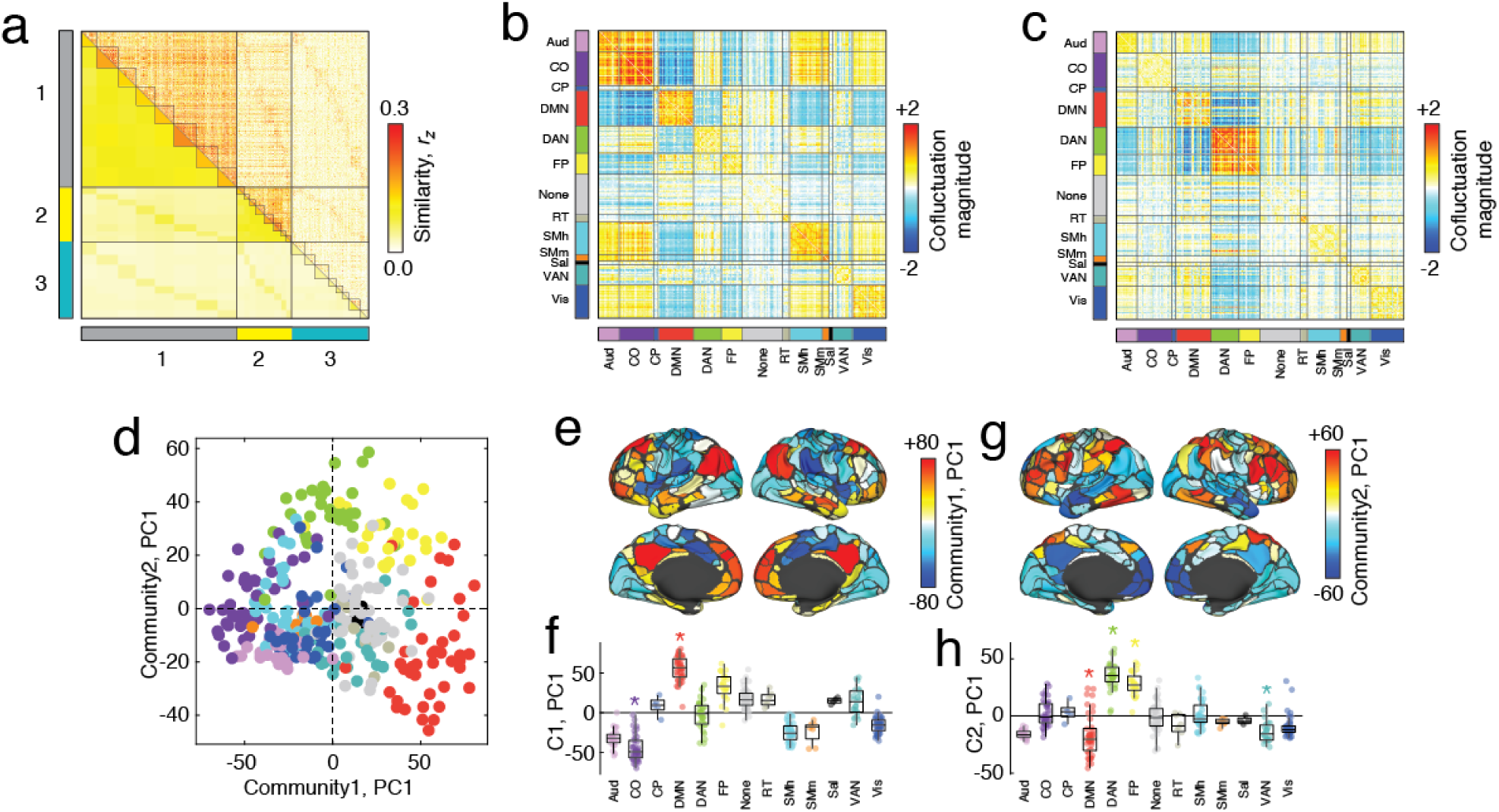
Clustering peak cofluctuation. (*a*) We clustered the cofluctuation similarity matrix using a multi-scale consensus clustering algorithm, resulting in two large communities (1 and 2) and a third set of much smaller communities, grouped together here to form community 3. Here, black lines divide communities from one another and, internally, participants from one another. The mean cofluctuation pattern for communities 1 and 2 are shown in *b* and *c*. To understand activity that underpins each community, we pooled together corresponding activity time series separately for communities 1 and 2 and performed principal component analysis on each set, returning the primary mode of activity (PC1). (*d*) Scatterplot of PC1 for community 1 and community 2. Colors denote brain systems. (*e*) Topographic depiction of PC1 for community 1. (*f*) PC1 grouped according to brain system. Asterisks indicate *p < p_adj_* (FDR fixed at *q* = 0.05). Panels *g* and *h* show corresponding plots for community 2. Analogous information about community 3 can be found in Fig. S7.

To better understand why certain cofluctuation patterns were grouped together, we examined group-representative centroids for each community by calculating the mean cofluctuation pattern of all frames assigned to that community (Fig. 5b,c). We found that community 1 reflected a topology that expressed strong and anticorrelated cofluctuations mostly between cinguloopercular and default mode networks (Fig. 5b). Community 2, on the other hand, expressed strong anticorrelations between the default mode with dorsal attention and fronto-parietal networks (Fig. 5b). See Fig. S9 for topographic depiction of systems on cortical surface.

We next wanted to better understand how brain *activity* drives the cofluctuation patterns described above. To do this, we extracted activity profiles (regional BOLD activity, rather than cofluctuations) during event peaks, grouped them by community, and performed principal components analysis (PCA). The first principal component for each community represents the mode of fMRI BOLD activity that tended to occur during frames assigned to that community. We found the first principal components of communities 1 and 2 (accounting for 25% and 26% of variance, respectively, with a sharp drop-off for increasing component numbers, Fig. S10) to be uncorrelated (*r* = − 0.025; *p* = 0.65; Fig. 5d). The first principal component for community 1 (Fig. 5e) exhibited significant activation of the default mode and inactivation of cingulo-opercular, visual, and somatomotor networks (distance-preserving permutation test of node order, i.e. spin test [36–38]; false discovery rate fixed at 5%; *p*_*adj*_ = 5.3 × 10^*−*4^; Fig. 5f). The first principal component for community 2 (Fig. 5h) exhibited significant activation of dorsal attention and fronto-parietal networks and inactivation of default mode, ventral attention, and visual networks (spin test; false discovery rate fixed at 5%; *p*_*adj*_ = 0.02; Fig. 5f). We show individual-level principal components in the **Supplementary Material** (Fig. S11) and derive similar modes of activity using an alternative procedure (Fig. S12). Note that the PCA analysis results in modes of activity; in all cases, the sign of PCs can be flipped and result in the same pattern of co-activity.

In the **Supplementary Material** we perform a similar analysis of Midnight Scan Club data parcels fit to each participant individually. We show that the first principal component of brain activity during high-amplitude co-fluctuations is similar irrespective of whether we use group-level or individualized parcels (Fig. S13). Even when we perform event detection separately using the individualized parcels, we find similar modes of activity (mean similarity of *r* = 0.89; *p <* 10^*−*11^; Fig. S14). Note that because the number of parcels differ across individuals, this comparison was carried out at a system level.

Again, we use MyConnectome data as a replication dataset, finding analogous communities (Fig. S15). We also take advantage of the fact that MyConnectome data includes many more samples from an individual than the Midnight Scan Club (84 scans versus 10), to identify several communities not evident in the group analysis of the Might Scan Club data. We also find evidence of interdigitated communities with similar system-level profiles but drawing on different regions from within those systems (Fig. S16).

These observations build on our previous study, which focused on a single pattern of cofluctuation during high-amplitude frames. Here, we use data-driven methods to show that high-amplitude cofluctuation is not monolithic and can be divided into meaningful sub-patterns, each driven by a distinct mode of brain activity.

### Event communities are individualized

In the previous section, we showed that cofluctuation patterns expressed at the peaks of high-amplitude events could be grouped into meaningful communities. Within each community, are these patterns individualized or are they shared across participants? Which node pairs are most variable between individuals and, therefore, more likely to be useful for subject fingerprinting and identifiability?

To address this question, we analyzed communities 1 and 2 separately and in greater detail. Although collectively each community is cohesive (similarity is greater among cofluctuation patterns assigned to the same community than to other communities; see Fig. S17 and Fig. S18), we also found evidence that the similarity between cofluctuation patterns is stronger still when they come from the same participant (t-test comparing within- and between-individual similarity; *p <* 10^*−*15^; Fig. 6a-d). Further, we identified the pairs of brain regions whose cofluctuations were most variable across individuals by computing the standard deviation of edge weights across participant centroids (Fig. 6e-f). Our rationale for doing so was that pairs of regions whose cofluctuation ampli-tude was variable are also among those most likely to drive individualization [39]. For the cofluctuation pattern expressed by community 1, we found that the most variable edges linked the cingulo-opercular network to the dorsal attention, fronto-parietal, and ventral attention networks (Fig. 6e). In the case of community 2, the most variable edges were linked to default mode, dorsal attention, and fronto-parietal.

**FIG. 6.**
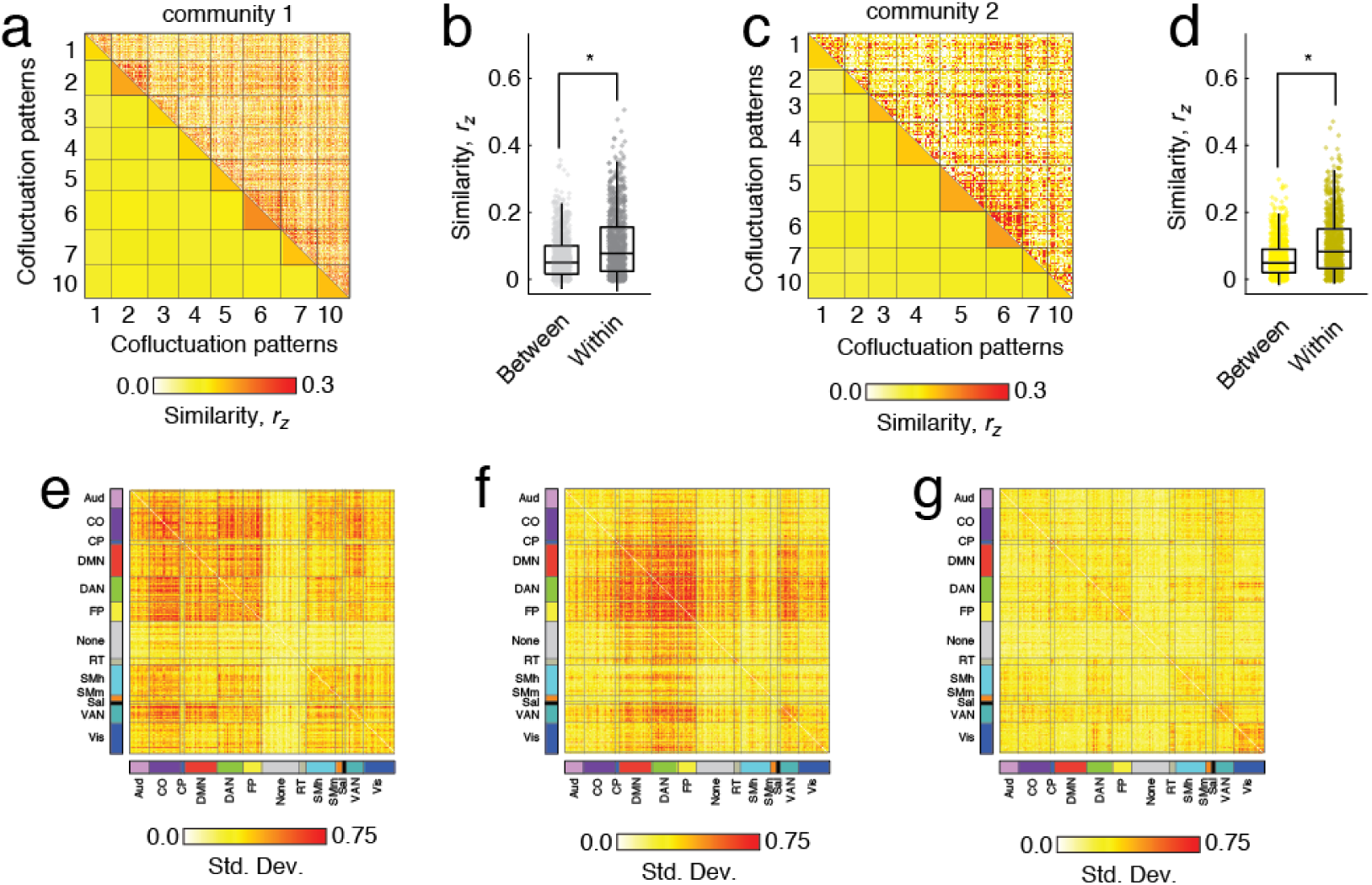
Personalization of cofluctuation patterns. We separately extracted cofluctuation patterns for communities 1 and 2 and computed the pairwise similarity. Similarity matrices are shown in panels *a* and *c*. In both cases, we found that withinindividual similarity was statistically greater than between-individual similarity. Boxplots of similarity scores are depicted in panels *b* and *d*. Asterisks indicate *p < p_adj_* based on permutation test (FDR fixed at *q* = 0.05). For each community, we calculated mean cofluctuation patterns for each participant and, across participants, computed the standard deviation of each node-pairs cofluctuation magnitude. The patterns of variability for communities 1, 2, and 3 are depicted in panels *e, f*, and *g*, respectively.

Note that in Fig. S19 and Fig. S20, we further explore the individualization of high-amplitude cofluctuation patterns. Briefly, we follow [40] and compute the similarity (correlation) of regional cofluctuation patterns estimated at the group and subject levels for both communities 1 and 2 (Fig. S19a,b). This procedure yields a similarity map for each subject and community (Fig. S19c-f), which we analyzed further. Specifically, we calculate the mean similarity of regions in each putative brain system [32] and demonstrated that, for community 1 regions in the ventral attention system was more dissimilar from the group than expected (space-preserving permutation test, 10000 repetitions, *p <* 10^*−*4^; Fig. S19f). For community 2, regions in the somatomotor-hand system, along with those that lack a clear system assignment, were more dissimilar. Interestingly, regions in the default mode and cingulo-opercular systems (community 1) and default mode, fronto-parietal, and dorsal attention systems (community 2) were more similar than expected. These systems all participate in the strongest co-fluctuations and largely typify each cluster. Finally, we investigate whether subject deviations from the group are evident in individual scan sessions. To test this, we repeat the above procedure but using scan-resolved estimates of communities (see Fig. S20 for an example from MSC06 and community 1). In general, we find that similarity maps for any given subject are highly repeatable across scan sessions and dissimilar between individuals (*t*-test of mean within- and between-subject similarity; *p <* 10^*−*15^; Fig. S19g).

These observations suggest that high-amplitude cofluctuation patterns reflect a topology that is broadly shared across individuals but is systematically and individually altered.

### Functional connectivity is modeled accurately only when using individual-specific co-fluctuation patterns

Previous studies have shown that time-averaged FC is individualized and can serve as a fingerprint of an individual [8, 12, 41]. Here and in [19], we show that time-averaged FC can also be approximated using only a small number of high-amplitude frames (events) and that the cofluctuation patterns expressed during those events can be grouped into a small number of clusters or communities. How do these different communities produce individualized patterns of FC? Are the cofluctuation patterns fixed at the group level but expressed in different proportions from one individual to the next? Or are the proportions fixed while the cofluctuation patterns vary idiosyncratically? Here, we present a model to adjudicate between these and related hypotheses.

Motivated by previous studies showing that FC can be described using only high-amplitude frames [3, 4, 6, 7, 19], our model assumes that FC depends exclusively on the cofluctuation patterns expressed during events and that low- and middle-amplitude frames make no contribution (Fig. 7a). Specifically, we model static FC as a linear combination of the centroids for communities 1, 2, and 3, weighted by how frequently those communities appear in the data. To fit this model, we must estimate centroids and frequencies. These estimates can be carried out separately for each subject, yielding subject-specific centroids and frequencies. However, estimates can also be made at the group level, yielding a set of centroids and frequencies that are shared across individuals. In both cases, we define model fit as the correlation of edge weights in the observed and predicted FC matrices.

**FIG. 7.**
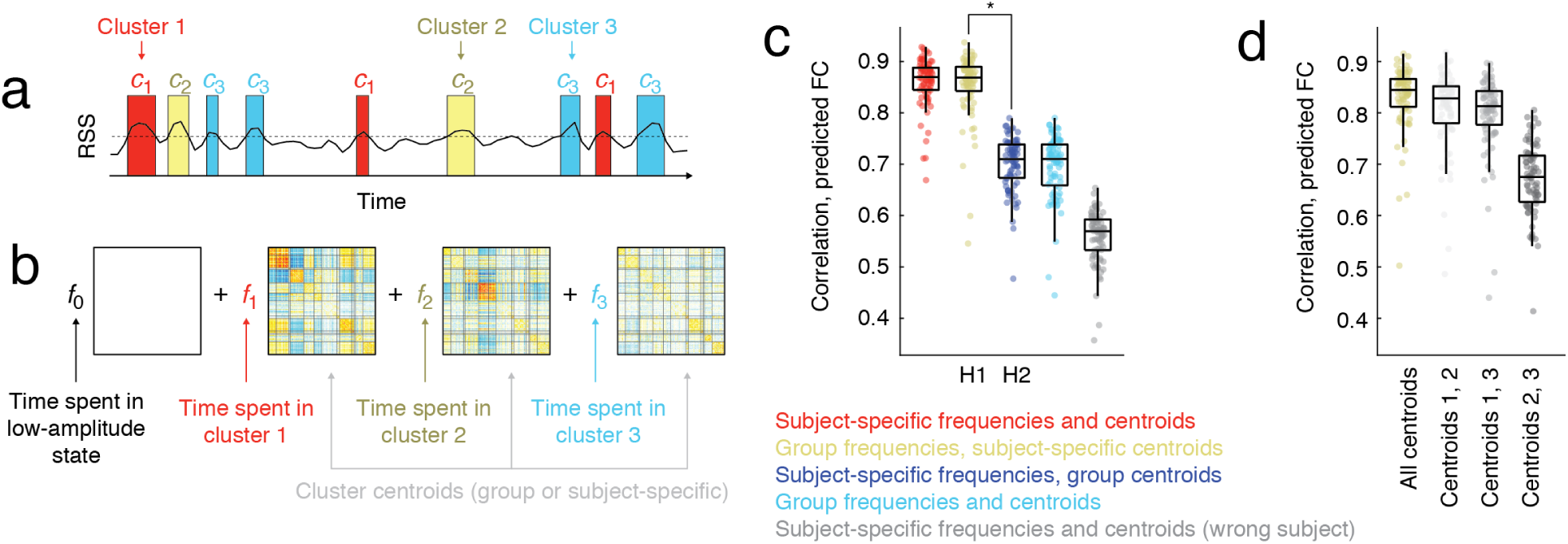
Modeling participant- and scan-specific FC with cofluctuation patterns. (*a*) We hypothesized that FC is driven by brief high-amplitude patterns of cofluctuation. A scan session can be abstracted, then, as periods of time where cofluctuation is close to zero punctuated by periods of time where cofluctuation corresponds to one of the cofluctuation communities. (*b*) We modeled FC as a linear combination of cofluctuation patterns corresponding to communities 1, 2, and 3 as well as a blank state where cofluctuation was treated as zeros (low-amplitude frames). We varied whether cofluctuation patterns and the frequencies with which they appear are estimated at the subject or group level. We also tested a model that used subject-level estimates of centroids and frequencies from other subjects. (*c*) Performance of the five models. Overall, models that included participant-specific information about cluster centroids outperformed other models. (*d*) We performed a sensitivity analysis of model 2, which used subject-specific centroids and group-level frequencies to predict FC patterns. In this analysis, we selectively exclude each of the three communities from the model so that it does not contribute to the prediction. We find that removing community 1 yields the biggest decrease in model performance, suggesting that it drives the model performance more so than communities 2 or 3.

Here, we test five model variants that combine subject- and group-level information in different configurations. The first two models make predictions of FC using subject-specific estimates of centroids and frequency estimates made at either the subject (model 1) or group level (model 2). Similarly, the next two models pair group-level estimates of centroids with either subject- or group-level estimates of frequencies (models 3 and 4). Finally, we test a fifth model that makes predictions of subject *s*’s FC using subject-specific centroids and frequencies estimated for the remaining seven subjects (different from the subject whose FC we are trying to predict), yielding seven independent predictions. We define this model’s fitness as the best prediction out of the seven.

Importantly, these models allow us to directly test the competing hypotheses that individualized FC is driven by subject-specific co-fluctuation patterns or subject-specific frequencies (labeled H1 and H2 in Fig. 1). Here, we found that model performance under H1 was significantly greater than performance for H2 (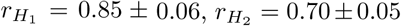; paired sample *t*-test, *p <* 10^−15^). Additionally, combining subject-specific centroids and frequencies yielded a small but statistically significant improvement in performance, largely by reducing the number of outlying points (*r* = 0.86 ± 0.05; paired sample *t*-test, *p* = 0.02). Collectively, these findings suggest that the subject-specificity of high-amplitude cofluctuation patterns drive the organization of static FC. Finally, we perform a sensitivity analysis to identify which of the three communities drive these effects. For all models, we find that model performance suffers the most by removing community 1, which alone accounts for 54.1% of all events (Fig. 7d).

Additionally, we replicated these findings using: MSC data processed using an alternative processing pipeline and parcellation [42, 43] (Fig. S21), at different spatial scales (100-node parcellation) [43] (Fig. S22), and without including global signal regression in the processing pipeline (Fig. S23). One notable discrepancy, however, was that in the absence of global signal regression, high-amplitude co-fluctuation patterns did not exhibit anticorrelations. This is in line with previous observations concerning the effect of the global signal on FC [44, 45]. Rather, the groups of brain regions that had previously engaged in anticorrelated behavior now formed their own distinct community, yielding four communities instead of two (along with a much smaller fifth community). As a consequence, we modeled FC as a linear combination of five states.

These results suggest that the cofluctuation patterns expressed at the peaks of high-amplitude events can, on their own, explain a significant fraction of participant-specific variance in time-averaged FC. More importantly, our results reaffirm that cofluctuation patterns during event peaks are participant specific; even when predicting a held-out scan, when centroids are estimated using participant-specific data, we find marked improvement in model performance.

## DISCUSSION

Here, we extended our recent analyses of edge time series and putative high-amplitude cofluctuation events [19, 26]. We proposed a simple null model that allowed us to identify frames whose amplitude was significantly greater or less than chance. We then analyzed the cofluctuation patterns expressed during these frames, discovering that across scans of the same individual, the cofluctuation patterns expressed during frames with lower-than-expected amplitude were dissimilar. In contrast, we found that the cofluctuation during high-amplitude frames was repeatable within an individual and dissimilar between individuals. We then clustered patterns of cofluctuation expressed during high-amplitude frames, identifying a small number of cofluctuation patterns that were shared across individuals. These patterns, however, were altered subtly yet systematically, so that they could be used to reliably distinguish participants from one another. Finally, we tested the hypothesis that FC could be predicted exclusively from the co-fluctuation patterns expressed during events, and constructed a model that generated estimates of FC given a set of cofluctuation community centroids and the frequency that those centroids are expressed by that individual. We found that the model performed well only when the centroids were estimated using data from the participants whose FC we were aiming to predict.

### High-amplitude cofluctuations can be partitioned into different communities based on their topology

Intrinsic or resting-state functional connectivity reflects the coupling of spontaneous activity between distant brain regions [1, 46]. It is often used to construct a graphical representation of the brain to be analyzed using tools from network science [47, 48]. Although inter-individual differences in FC have been linked to an individual’s clinical [16], cognitive [49], and developmental state [15], the dynamic origins of individualized FC remain unknown.

Recently, we presented a method for decomposing FC into its framewise (instantaneous) contributions [19, 23]. Our work, in agreement with other recent studies [3, 4, 6, 7], demonstrated that all frames do not contribute equally to FC – rather only a small number of high-amplitude frames – “events” – when averaged together, are necessary for explaining a high proportion of variance in FC. In that study, however, we only examined the mean pattern of high-amplitude cofluctuations and did not investigate individual events nor did we characterize variation in the co-fluctuation patterns across events.

Here, we address these issues using two dense phenotyping datasets. Leveraging a statistical test for identifying high-amplitude frames, we show that “events” are *not* monolithic and comprise distinct patterns of cofluctuations. Using a data-driven clustering method, we find evidence of two patterns of cofluctuation that are conserved across all participants and scans. These patterns emphasize opposed activation of default mode regions with cingulo-opercular and sensorimotor systems (community 1) and with dorsal attention and fronto-parietal systems (community 2). We also find evidence of smaller communities corresponding to less frequent events involving only a fraction of participants. In the main text we grouped these patterns into a single community, but find that the two largest (accounting for 8.6% of event peaks) involve cofluctuations of sensorimotor systems, which we explore in the **Supplementary Material**.

The two largest communities have interesting properties. For instance, the default mode is cohesive (densely interconnected, internally) in both, but is selectively decoupled from distinct sets of brain systems associated with processing sensorimotor information [50] and coordinating flexible, goal-directed behavior [51]. In neither community does the default mode couple strongly to other systems. Rather, in these high-amplitude states it maintains relative autonomy, in agreement with studies that have examined its “hubness” using the participation coefficient [52] – a graph-theoretic measure that describes the extent to which a node’s connections are distributed across or concentrated within communities [53].

Another interesting feature of these communities is their possible relationship to network states obtained by clustering sliding-window estimates of time-varying FC [54] or co-activation patterns (CAPs) [55]. Indeed, these approaches all yield estimates of repeating patterns of co-activity and connectivity. Although recent work suggests a deep mathematical relationship among these approaches and static FC, the precise nature of this link remains undisclosed [56]. Here, we analyze edge time series, a parameter-free method for estimating instantaneous cofluctuations between regional activity (localized to individual frames) and whose sum is precisely time-averaged FC. In contrast, sliding-window estimates of time-varying FC require users to specify a window duration and overlap fraction (the number of frames shared by successive estimates of FC) and, due to the sliding window, lead to temporally blurred connectivity estimates that cannot be precisely localized in time [57, 58]. In fact, we speculate that brief high-amplitude events may be present in sliding-window estimates of time-varying FC, but because they evolve over timescales much shorter than that of the typical window duration, are effectively obscured due to blurring. CAPs, on the other hand, which leverages a similar procedure as ours, identifies repeated and high-amplitude patterns of activity rather than cofluctuation matrices (the product of instantaneous activity patterns). For this reason, CAPs traditionally does not offer the mathematical link between cofluctuations and FC that edge time series does [59]. Critically, then, because edge time series are a decom-position of FC, events can be viewed as the “atoms” or “building blocks” of FC.

Collectively, our findings suggest that cofluctuation patterns expressed during putative network-wide events are variable but can be described in terms of two principal patterns. These findings extend our previous study [19] and open up opportunities for future studies to investigate inter-individual differences in these patterns as well as the smaller and less frequent patterns.

### High-amplitude cofluctuation patterns are individualized and drive time-averaged FC

One of the questions we aimed to address was whether the individualization of FC occurred because: *a*) high-amplitude cofluctuation patterns are shared across individuals but expressed in different subject-specific proportions or *b*) cofluctuation patterns expressed during high-amplitude events are inherently subject-specific. The answer to this question is important, as it speaks to the origins of individual differences in FC [22, 40], has implications for brain-behavior studies [14], and also informs our understanding of time-varying FC [60].

Here, we addressed these questions by aggregating and clustering high-amplitude cofluctuation patterns from across all participants. This analysis returned two large communities in which every participant and scan were represented, indicating that, to some extent, patterns of high-amplitude cofluctuations are indeed shared across individuals. However, when we examined these communities in greater detail, we found that within communities there existed more cohesive sub-communities corresponding to individual participants.

To better adjudicate between hypotheses, we constructed a simple model to predict an individual’s scan-specific pattern of FC. Motivated by previous studies [3, 4, 6, 7], this model assumed that FC is driven exclusively by high-amplitude cofluctuations and that all other time points made negligible contributions to FC. We then replaced FC during high-amplitude frames with the centroid of the community to which those frames were assigned. We found that, when centroids were generated using data pooled from across all participants we could explain only ≈ 50% of the variance in functional connection weights. However, when we estimated centroids using data from the same participant whose FC we were predicting, the model exhibited a significant increase in performance, accounting for ≈ 75% of variance in connection weights.

These observations suggest that incorrectly ascribing group-level features to an individual participant distorts our prediction of their FC, favoring the hypothesis that high-amplitude cofluctuation patterns are individ-ualized. This endorsement of hypothesis *b* comes with some caveats, however. Although FC is the product of individualized cofluctuation patterns, those patterns appear to be variants of archetypal patterns, i.e. the two large communities discussed in the previous section. These observations align with other studies showing that the individualization of FC is generally a subtle modulation of features that are evident in group-level data, from brain systems [61–63] to regional FC fingerprints [8]. Notably, these findings also corroborate other “edge-centric” analyses of individualized patterns of brain connectivity. For instance, in [64], the authors demonstrated that the correlation structure of edge time series – a construct referred to as “edge functional connectivity”-outperformed traditional FC in terms of identifiability. Moreover, reconstructions of eFC using principal components further improved its performance. Those results, paired with those of the present study, suggest that edge-based approaches may offer a useful framework for investigating individualization of connectivity and, eventually, linking it back to behavioral, cognitive, and clinical phenotypes.

Our results have implications for studies of brain-behavior correlations as well as state-based analyses of time-varying FC. We show that inter-individual differences in FC are largely shaped by differences in high-amplitude cofluctuation patterns. However, our sensitivity analysis (Fig. 7d) demonstrated that of the three communities we considered, one contributed dispropor-tionately to the individualization of FC relative to the other two. This suggests that, rather than linking inter-individual differences in FC across individuals, it may be more profitable to directly investigate specific community centroids, e.g. those that drive individual variation, potentially leading to improvements in brain-behavior correlations.

Our results also have implications for studies of network states in time-varying FC [54, 58]. In general, these studies cluster time points together based on the similarity of networks to one another. To facilitate ease of comparison across individuals, this step is usually performed using concatenated data from many participants or conditions. Different metrics can be calculated from these partitions, e.g. cumulative time a participant spends in any cluster, transition matrices, etc., and linked to behavioral and clinical phenotypes. Our results suggest that, although methodologically convenient, clustering time points together and treating them as recurrences of the same “network state” likely obscures meaningful participant-level variation.

### Future directions

The results presented here raise important questions that should be investigated in future research. First, because events contribute disproportionately to the organization of time-averaged FC and because they appear to be drivers of individualization, they should be the target of future studies. We investigated events in two dense sampling studies and in a total of nine brains. Although these data allowed us to investigate the extent to which cofluctuation patterns during events are shared *versus* individualized, the small number of participants precludes the possibility of investigating behavioral, cognitive, or disease correlates of events [65]. Future studies should investigate communities of high-amplitude events in larger datasets.

Relatedly, our study examined event structure exclusively during task-free resting-state conditions. We demonstrated that time-averaged FC could be well-approximated using only high-amplitude frames and individualized estimates of cofluctuations during those frames. What happens to events when participants are asked to perform tasks in the scanner? Previous studies have demonstrated that tasks systematically modulates patterns of FC [49]. Do these changes reflect different patterns of cofluctuations during events? Are they the same patterns as rest but in different proportions? And directly related to the aims of this study, are task events similarly personalized or can they be used to strengthen brain-behavior associations [66]?

Here, we focused on the contributions of high-amplitude “events” to patterns of time-averaged FC. The simple model we proposed even goes so far as to consider contributions from all other frames as negligible. Is this really the case? What biases might we reinforce by focusing on high-amplitude frames? High-amplitude cofluctuations make proportionally bigger contributions to time-averaged FC than low-amplitude cofluctuations. This statement is non-controversial; edge time series are a mathematically precise “temporal unwrapping” of the Pearson correlation into its framewise contributions, the average of which is simply FC [19, 23–26]. For this reason, it makes sense to focus on frames where many edges simultaneously make big contributions – those same frames necessarily will, on average, make bigger contributions to FC than, say, frames where only a few edges exhibit high-amplitude edge time series. However, this does not rule out the possibility that frames outside of high-RSS events, which are more numerous, make contributions that outweigh or match those of high-amplitude frames. Additionally, in focusing on *global* high-amplitude events, we may miss out on events involving small brain systems, which will fail to meet statistical criteria for significance due to their size.

Edge time series represent only the latest in a series of methods for tracking and modeling time-varying changes in networks that include time-frequency analysis [67], sliding windows [57], instantaneous phase-locking [68], co-activation patterns [55, 69], multiplication of temporal derivatives [70], quasi-periodic patterns [71], and model-based frameworks [72]. Broadly, these different approaches can be classified on the timescales over which they describe changes in brain activity or connectivity. Some, like CAPs, edge time series, and instantaneous phase-locking, characterize changes at a frame-wise timescale. Others require windowing or smoothing of data to track time-varying fluctuations. Of particular interest are quasi-period patterns, (QPPs), which refer to repeated sequences of brain activity [73–75]. Recently, [76] developed a method for assessing contributions to FC by repeating patterns. In essence, they “regress out” each QPP’s time course from the BOLD signal and calculate FC before and after doing so, allowing the authors to assess the contribution of the QPP to FC. More recently, [77] identified three QPPs and demonstrated that, using only the time courses of those QPPs, could reconstruct and FC matrix that was correlated with the FC matrix estimated using the entire BOLD signal. Although these studies collective suggest that QPPs contribute to the overall pattern of FC, the precise mathematical link remains inexact. Moreover, every QPP evolves over an extended period of time and includes multiple frames, further complicating its relationship with FC. In contrast, edge time series are an exact decomposition of FC into its framewise contributions. That is, the mean of a given edge time series is mathematically equivalent to that edge’s weight in the FC matrix. This makes it possible to assess the collective contribution of individual frames to the overall FC pattern. In contrast, with QPPs and co-activation patterns (CAPs), the mathematical link to FC is not precise. Irrespective of whether their contributions can be determined precisely, QPPs, CAPs, iCAPs, “events” and related methods make clear that variation in activity and co-activity across time contribute to FC. Future work should focus not only on assessing the relative strengths and weakness of these methods but identifying underlying structural similarities between approaches [56].

Here, we analyze two dense-sampling datasets in which a small number of individuals were scanned many times. Our work is naturally extended by examining inter-individual variation of high-amplitude co-fluctuation patterns and community frequency in large, cross-sectional datasets, e.g. the Human Connectome Project [78]. In addition to imaging data, the HCP dataset includes rich cognitive, behavioral, and clinical assessments of participants. Future work should focus on linking inter-individual variation in those measures with high-amplitude events.

Finally, a key overarching and open question concerns the origins of high-amplitude cofluctuations. In our previous study, we demonstrated that movie-watching leads to synchronization of events across participants [19], suggesting that their timing can be modulated selectively, in that case by sensory input. But what about rest? In that previous study, we found no differences in event amplitude between rest and movie-watching, suggesting that spontaneous events are just as large as those driven by sensory input. A couple recent studies help us speculate on the origins of events. One possibility is that events help preserve brain circuit function in the absence of use. In [79], the authors demonstrated that disuse of motor circuits by casting participants’ arms leads to increases in high-amplitude pulses and that manually censoring these pulses reduces FC magnitude [80]. Another possibility is that burst activation of distinct systems, including default mode networks, co-occurs with the recall or previously-observed stimuli [81]. Yet another possibility, and one that is supported by recent results from electrophysiological studies [82], is that rapid and antagonistic (anti-correlated) relationships between specific subnetworks, including the default mode, are strongly associated with sustained attention, suggesting physiological origins. In all cases, these activation patterns are likely constrained by the underlying anatomical connectivity and reflect groups of mutually connected brain regions [30, 83]. A final possibility is that events are truly stochastic and are mere bi-products of modular correlated time series, where the activation of one element within a module implies the activation of the others [56]. In any case, future experimental studies – possibly invasive studies that allow for more targeted and temporally resolved recording – should be directed to investigate their origins.

## Limitations

The conclusions of this study are limited in several ways. Notably, we analyze fMRI data, which affords whole-brain coverage but at a spatial resolution of millimeters and a temporal resolution of, at best tenths of a second. Moreover, fMRI BOLD samples a slow signal slowly, and is only indirectly related to population activity. However, many studies have demonstrated a correspondence between BOLD fluctuations and FC estimated from fMRI with other modalities, including local field potentials [84], intracranial EEG [85], and optically recorded calcium imaging signals [86], positing a neural basis for functional connectivity estimated from the fMRI BOLD signal. Future studies could apply methods similar to those used here to other imaging modalities.

A second limitation concerns data quality, in-scanner motion, and other artifacts, which are known to impact estimates of FC and can produce burst-like behavior in fMRI time series [87, 88]. Here, we adopted a conservative approach and discarded putative events that occurred near censored frames. Notably, this procedure impacted low-amplitude frames to a greater extent than high-amplitude frames, suggesting that high-amplitude frames, in addition to contributing disproportionately to FC, are also less likely to be contaminated by artifacts. Nonetheless, how to adequately address motion-related issues in the analysis of FC remains an ongoing and disputed topic [89–91].

A final limitation concerns the use of PCA to extract “modes” of activity underlying communities. While we find that each community’s first component explains ≈ 25% variance, we do not investigate the remaining components, which likely include other meaningful modes of activity. Future studies should further investigate the link between brain activity and connectivity. Because edge time series is a mathematically exact decomposition of FC into its time-varying contributions, it represents a useful framework for doing so.

## Conclusion

In conclusion, we find that FC can be explained using a small number of high-amplitude frames. These frames can be clustered into a small number of communities corresponding to archetypal patterns of cofluctuation, broadly shared across individuals. However, these patterns undergo refinement at the level of individual participants, yielding reliable and individualized cofluctuation patterns. Finally, we show that participants’ FC is more accurately predicted using participant-specific estimates of their high-amplitude cofluctuation patterns compared to group-level estimates. Our study discloses high-amplitude, network-wide cofluctuations as dynamical drivers of individualized FC and introduces methodology for exploring their role in cognition, development, and disease in future studies.

## MATERIALS AND METHODS

### Datasets

#### Midnight Scan Club

The description of the Midnight Scan Club dataset acquisition, pre-processing, and network modeling is described in detail in [11]. Here, we provide a high-level overview. Data were collected from ten healthy, righthanded, young adult participants (5 females; age: 24-34). Participants were recruited from the Washington University community. Informed consent was obtained from all participants. The study was approved by the Washington University School of Medicine Human Studies Committee and Institutional Review Board. This dataset was previously reported in [11, 12] and is publicly available at https://openneuro.org/datasets/ds000224/versions/00002. Imaging for each participant was performed on a Siemens TRIO 3T MRI scanner over the course of 12 sessions conducted on separate days, each beginning at midnight. In total, four T1-weighted images, four T2-weighted images, and 5 hours of restingstate BOLD fMRI were collected from each participant. For further details regarding data acquisition parameters, see [11].

High-resolution structural MRI data were averaged together, and the average T1 images were used to generate hand-edited cortical surfaces using Freesurfer [92]. The resulting surfaces were registered into fs LR 32k surface space as described in [93]. Separately, an average native T1-to-Talaraich [94] volumetric atlas transform was calculated. That transform was applied to the fs LR 32k surfaces to put them into Talaraich volumetric space.

Volumetric fMRI pre-processing included including slice-timing correction, frame-to-frame alignment to correct for motion, intensity normalization to mode 1000, registration to the T2 image (which was registered to the high-resolution T1 anatomical image, which in turn had been previously registered to the template space), and distortion correction [11]. Registration, atlas transformation, resampling to 3 mm isotropic resolution, and distortion correction were all combined and applied in a single transformation step [95]. Subsequent steps were all completed on the atlas transformed and resampled data. Several connectivity-specific steps were included (see [96]): (1) demeaning and de-trending of the data, (2) nuisance regression of signals from white matter, cerebrospinal fluid, and the global signal, (3) removal of high motion frames (with framewise displacement (FD) > 0.2 mm; see [11]) and their interpolation using power-spectral matched data, and (4) bandpass filtering (0.009 Hz to 0.08 Hz). Functional data were sampled to the cortical surface and smoothed (Gaussian kernel, *σ* = 2.55 mm) with 2-D geodesic smoothing.

The following steps were also undertaken to reduce contributions from non-neuronal sources [96, 97]. First, motion-contaminated frames were flagged. Two partic-ipants (MSC03 and MSC10) had high-frequency artifacts in the motion estimates calculated in the phase encode (anterior-posterior) direction. Motion estimate time courses were filtered in this direction to retain effects occurring below 0.1 Hz. Motion contaminated volumes were then identified by frame-by-frame displacement (FD, described in [98]), calculated as the sum of absolute values of the differentials of the 3 translational motion parameters (including one filtered parameter) and 3 rotational motion parameters. Frames with FD > 0.2 mm were flagged as motion-contaminated. Across all participants, these masks censored 28%±18% (range: 6% - 67%) of the data; on average, participants retained 5929±1508 volumes (range: 2733 – 7667). Note that in this paradigm, even the worst participant retained almost two hours of data. Nonetheless, we excluded two subjects from all analyses, both of whom had fewer than 50% usable frames in at least five scan sessions (MSC08 in 7/10 and MSC9 in 5/10). See Fig. S24 for a summary of usable frames for each subject and scan.

Time courses were extracted from *N* = 333 cortical regions using a common (group) functional parcellation [32]. We also analyze time courses estimated from using individualized parcellations (see [41] for details). Both group and individualized time series were used for FC estimation and edge time series generation.

#### MyConnectome dataset

All data and cortical surface files are freely available and were obtained from the *MyConnectome Project*’s data-sharing webpage (http://myconnectome.org/wp/data-sharing/). Specifically, we studied pre-processed parcel fMRI time series for scan sessions 14–104. Details of the pre-processing procedure have been described elsewhere [10, 31]. Each session consisted of 518 time points during which the average fMRI BOLD signal was measured for *N* = 630 parcels or regions of interest (ROIs). With a TR of 1.16 s, the analyzed segment of each session was approximately 10 minutes long.

### Functional connectivity

Functional connectivity (FC) measures the statistical dependence between the activity of distinct neural elements. In the modeling of macroscale brain networks with fMRI data, this usually means computing the Pearson correlation of brain regions’ activity time series. To calculate FC for regions *i* and *j*, then, we first standardize their time series and represent them as z-scores. We denote the z-scored time series of region *i* as **z**_*i*_ = [*z*_*i*_(1), …, *z*_*i*_(*T*)], where *T* is the number of samples. The Pearson correlation is then calculated as:

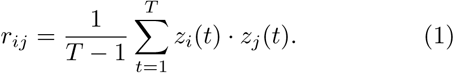

In other words, the correlation is equal to the temporal average of two regions’ cofluctuation.

### Edge time series

Recently, we proposed a method for decomposing FC into its framewise contributions. This is accomplished by simply omitting the averaging step in computing Pearson’s correlation. This omission results in a new time series:

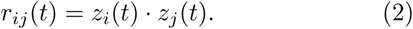

where the value of *r*_*ij*_(*t*) indexes the instantaneous cofluctuation between regions *i* and *j* at time *t*. When regions *i* and *j* both deflect from their mean in the same direction *r*_*ij*_ > 0, when they deflect in opposite direction, *r*_*ij*_ *<* 0, and when one (or both) of their activities is near their mean then *r*_*ij*_ ≈ 0. Importantly, the mean of this time series is exactly equal to the FC between regions *i* and *j*, and therefore we can think of *r*_*ij*_(*t*) as the instantaneous contribution of frame *t* to the overall FC.

If we consider the set of cofluctuation between all pairs of regions {*ij*} at time *t*, we can arrange those elements into a node-by-node connectivity matrix and analyze it as a network. We can also calculate the total amplitude of cofluctuation between all node pairs as their root sum square, 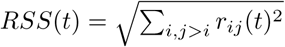.

### Multiresolution consensus clustering

We used a variation of modularity maximization [99] to group cofluctuation patterns into clusters or “communities”. Briefly, modularity maximization is a community detection method for partitioning relational data, e.g. networks, into non-overlapping communities. This is accomplished by optimizing a modularity quality function:

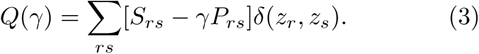

In this expression, *S*_*rs*_ is the similarity of cofluctuation patterns *r* and *s*; *P*_*rs*_ is the level of similarity expected by chance; *δ*(*z*_*r*_, *z*_*s*_) is the Kronecker delta function and is equal to 1 when the community assignments of patterns *r* and *s*, denoted as *z*_*r*_ and *z*_*s*_, are identical, and 0 otherwise. The structural resolution parameter, *γ*, controls the importance of *S*_*rs*_ relative to *P*_*rs*_ and, in effect, can be tuned to recover smaller or larger communities.

Here, we use modularity maximization to obtain a representative set of multi-resolution communities [34]. That is, a partition of cofluctuation patterns into commu-nities that takes into account how strongly coupled patterns are to one another at different scales, from finescale partitions of patterns into many small communities to coarse partitions of patterns into a few large communities. To do this, we sample communities at various scales by changing the value of *γ*. Specifically, we sample 10,000 different values of *γ* based from the distribution of *S*_*rs*_ values. At each value, we use a generalized version of the Louvain algorithm to optimize the corresponding *Q*(*γ*) [35] (http://netwiki.amath.unc.edu/GenLouvain/GenLouvain).

This procedure results in 10000 estimates of communities at a range of scales. We transform these estimates into a probabilistic co-assignment matrix, whose element *T*_*rs*_ is equal to the fraction of the 10000 partitions in which nodes *r* and *s* were assigned to the same community. From this matrix, we construct a new modularity:

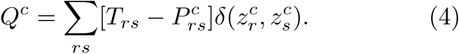

In this expression, 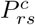 is the expected co-assignment of *r* and *s* to the same community and can be estimated as the mean value of the elements in the empirical coassignment matrix. 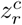 and 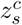 are the consensus commu-nity assignment of patterns *r* and *s*, respectively.

Optimizing *Q*^*c*^ tends to return partitions that are, as a group, more similar to one another and, possibly, identical. If this is the case, the algorithm ends and the resulting partition is accepted as the representative consensus partition. However, if there is any variability in the output so that the algorithm does not arrive at the same solution each run, the a new co-assignment matrix is estimated and its modularity optimized. This procedure repeats until the algorithm converges. We use the same generalized version of the Louvain algorithm to optimize the consensus modularity function and detect consensus communities.

### Principal component analysis of communities

The multiresolution consensus clustering procedure groups a series of vectorized *N* × *N* high-amplitude, cofluctuation matrices into communities. Each clustered cofluctuation matrix corresponds to the peak frame in a temporally contiguous series of frames whose RSS was significantly greater than that of a null model. While our primary aim was to understand how these matrices contribute to time-averaged FC, we were also interested in characterizing what types of activity, i.e. *N* × 1 patterns of fMRI BOLD data, give rise to high-amplitude cofluctuations.

Naively, one could address this question by identifying the frames corresponding to peak co-fluctuations and average over their corresponding activity patterns. This approach, however, can yield misleading results. This is because every co-fluctuation matrix can be generated by two different patterns of activity that are identical to one another except for their signs. To illustrate this, consider the toy case presented below. Suppose we have a four-node network with the following co-fluctuation matrix:

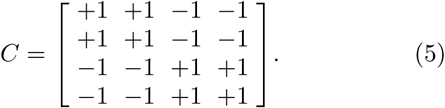

The elements of this cofluctuation matrix are given by *C*_*ij*_ = *z*_*i*_*z*_*j*_, where *z*_*i*_ is the z-scored activity of node *i*. Accordingly, this matrix could have been generated by either of the following patterns of activity:

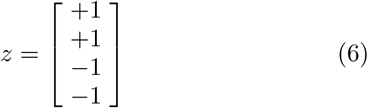

or

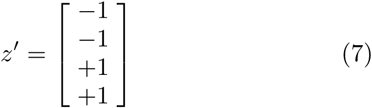

The simple average of *z* and *z*^*′*^ is a vector of zeros, which is unrelated to the co-fluctuation matrix. Averaging peak activity patterns can yield a similarly misleading result. Fortunately, both patterns are co-linear and satisfy the relationship *z* = − 1 · *z*′. That is, these patterns can be described by a single “mode” of activity, which we can detect by applying principal components analysis (PCA) to the activity patterns directly.

With this in mind, our strategy for characterizing the patterns of activity underpinning high-amplitude cofluctuations was as follows. For each of the cofluctuation matrices assigned to community 1, 2, and 3, we identified the scan and frame number in which they originated. Separately for each community, we aggregated the corresponding patterns of activity into matrices with dimensions of *N* × *N*_*peaks*_. Here, *N*_*peaks*_ is the number of high-amplitude frames assigned to a given community. Then, we applied PCA to these matrices, yielding a series of orthogonal components of dimension *N* × 1 and the variance explained by each component. Note that these components are used mostly for visualization and to better understand the link between cofluctuation matrices and brain activity.

In the main text, we also describe a second approach for uncovering the dominant mode of activity underpinning the centroids of communities. Each centroid represents the average over many cofluctuation matrices. To discover the optimal mode of activity, we aimed to determine the elements of *z* = [*z*_1_, …, *z*_*N*_] that minimized the following cost function:

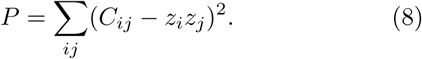

To optimize *P*, we used a greedy algorithm which we repeated 100 times. Briefly, we initialized the algorithm with a *z* ∈ *ℝ*^*N ×*1^ vector whose elements were drawn independently from 𝒩(0, 1) and calculated the corresponding cost of *P*. Then we randomly selected a node, *i*, replaced its current value with another value randomly drawn from the same distribution. We denote the resulting vector as *z*′ and its cost as *P*′ If *P*′ *< P* then we retained *z*′. We repeated this procedure 25000 times, gradually reducing *P*. In practice, we found that the algorithm converged to highly similar solutions (mean similarity across 100 runs of *r* = 0.993.

### Predictive model of FC

In the main text we described a procedure for modeling FC in terms of cofluctuation community centroids. In this section, we provide more details of how the model works.

In our previous work [19], we claimed that FC is driven by high-amplitude frames. One way to test whether this is the case is to “zero out” all low-amplitude and non-significant frames and to compute FC as the sum of whatever cofluctuation patterns are expressed at high-amplitude frames. Here, we take this claim one step further and state that the cofluctuation patterns expressed during high-amplitude frames are recurrences of one of three template patterns, which we obtained from the community detection analysis.

This intuition can be formalized by the following model:

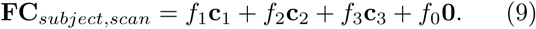

In this expression, *f*_1_, *f*_2_, and *f*_3_ are the fractions of all low-motion frames in which a participant expresses communities 1, 2, and 3, respectively. The parameter *f*_0_ is the fraction of frames in which a participant is in a low-amplitude or non-significant state. The values of these parameters come from the results of the multiresolution consensus clustering analysis. The other parameters **c**_1_, **c**_2_, and **c**_3_ represent the average pattern of cofluctuation for of the three communities. The final parameter, **0** is a node-by-node matrix where all elements are zero.

The first two models make predictions of FC using subject-specific estimates of centroids and frequency estimates made at the subject- and group-level (models 1 and 2). Similarly, the next two models pair group-level estimates of centroids with subject- and group-level estimates of frequencies (models 3 and 4). Finally, we test a fifth model that makes predictions of subject *s*’s FC using subject-specific centroids and frequencies estimated for the remaining seven subjects. For this model, we retain the best fit of the seven.

Note that when predicting FC for a given scan, both subject- and group-level centroids and frequencies are estimated while excluding observations from that scan. Note also that frequency is estimated as the total number of frames associated with a given community. To obtain this number, we first map the community assignments of event cofluctuation patterns back to the segment they originated (a temporally contiguous set of frames whose amplitude is significantly greater than that of the null model). We then assign the same community label to all frames that make up that segment. Finally, we calculate the frequency of each community as the total number of frames assigned to that community divided by the total possible number of frames.

In general, the model can generate matrices that are not positive semidefinite (all positive eigenvalues) and therefore not possible correlation matrices. Accordingly, we transform each matrix (translation/rotation/scaling) to match the nearest admissible matrix by minimizing the Frobenius norm using the MATLAB function nearcorr.m. In all cases, we measure model fitness as the correlation of upper triangle elements in the true FC with those of the predicted FC matrix.

## DATA AVAILABILITY

Midnight Scan Club raw data and derivatives are available here: https://openfmri.org/dataset/ds000224/ and in processed, parcellated form here https://www.dropbox.com/sh/tb694nmpu2lbpnc/AABKU_Mew7hyjtAC4ObzGVaKa?dl=0. Processed and parcellated MyConnectome data is available here: http://myconnectome.org/wp/data-sharing/.

## AUTHOR CONTRIBUTIONS

RFB designed study, analyzed data, wrote code, generated figures, and wrote initial draft of manuscript. SC, SG, and OS designed study, revised manuscript, and wrote final version of manuscript.

## ACKNOWLEDGMENTS

We thank Evan Gordon for sharing individualized parcellations for MSC participants. This material is based upon work supported by the National Science Foundation under Grant No. 076059-00003C (RFB, OS).

---

**FIG. S1.**
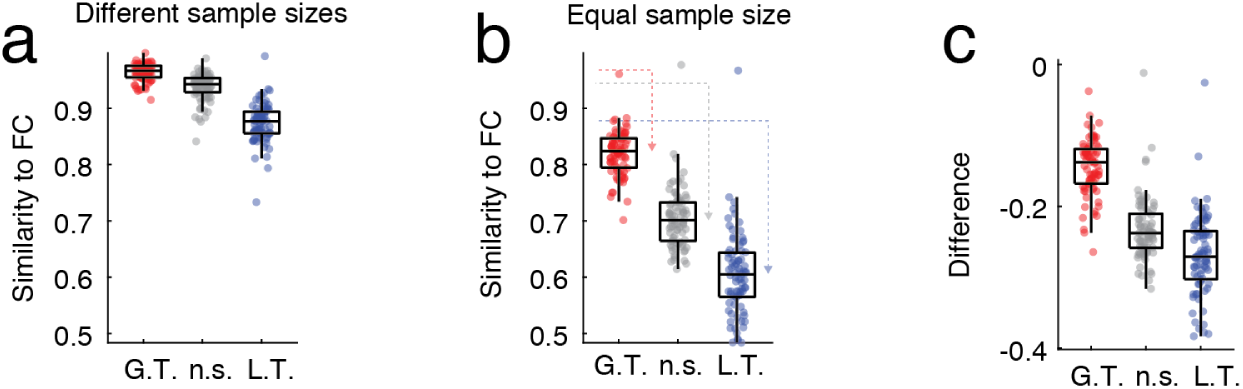
Controlling for sample size when comparing high-amplitude, non-significant, and low-amplitude frames with functional connectivity. In the main text, we demonstrated that FC is differentially correlated with reconstructions made using only high-amplitude, low-amplitude, or non-significant frames. In general, we found that high-amplitude frames outperformed the other two categories (see panel *a*). However, the correlation magnitude of FC with the three categories was greater than what had been previously reported [19]. The discrepancy is likely due to differences in the number and duration of frames used in reconstructing each matrix. Moreover, the number of frames differed systematically across categories. To address these concerns, we repeated the analysis reported in the main text using an equal number of randomly-sampled frames from each category (27 frames *≈* 1 minute; 100 repetitions). We find that the correlation of the reconstructions with FC decrease across all three categories (panel *b*) but that the decrease is steepest for the non-significant and low-amplitude frames, suggesting that their similarity to FC had been inflated due to the largest number of samples (panel *c*).

**FIG. S2.**
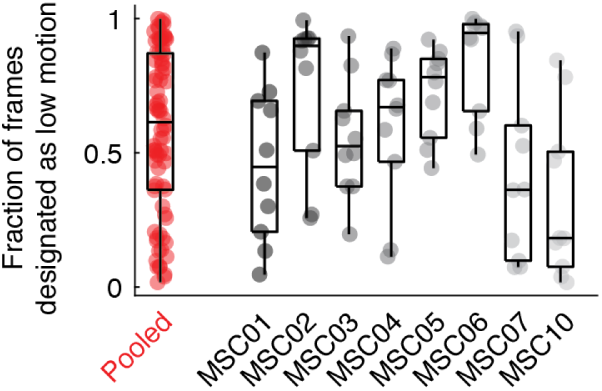
Variability in low-amplitude co-fluctuations reflects stable inter-individual differences in motion. In the main text, we find that low-amplitude frames pooled across subjects and scans are variable in terms of corresponding in-scanner motion measures. To better understand the origins of this variability, we separated the motion estimates by subject. We found that there was a significant difference across individuals in terms of motion (ANOVA, *F* (8) = 4.7, *p* = 0.0002). These findings suggest that some of the variation in the motion estimates for low-amplitude frames is attributable to inter-individual differences.

**FIG. S3.**
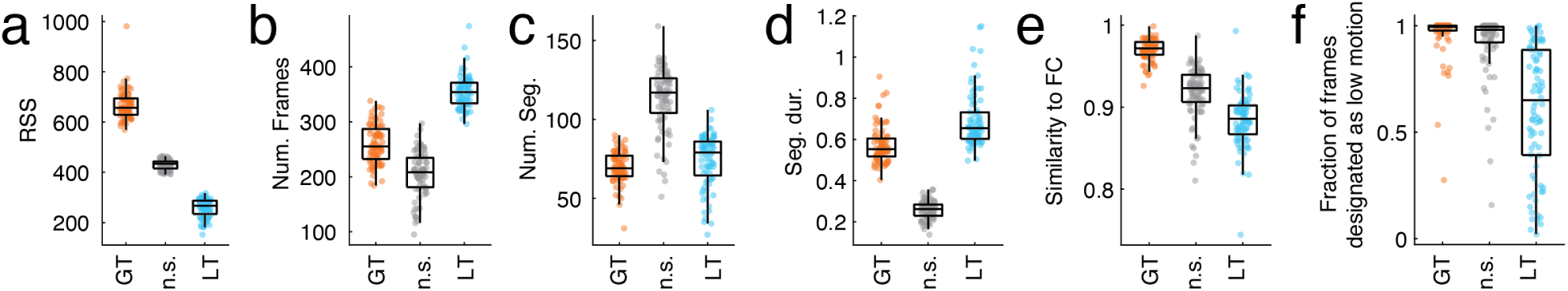
Statistical comparison of high- and low-amplitude frames using individualized parcels and Midnight Scan Club data. In the main text, we described a statistical procedure for partitioning frames into three categories based on how their RSS values compared to those of a null model. We applied that model to data from the Midnight Scan Club in which participants’ brains were parcellated into individualized regions and compared the three categories across multiple features, including their RSS amplitude, number of frames associated with each category, the number of contiguous frames of the same category (we call these “sequences”), sequence duration, the similarity of FC reconstructed using frames of each category to the static FC matrix, and the fraction of frames of a given category that were censored due to motion/data quality issues.

**FIG. S4.**
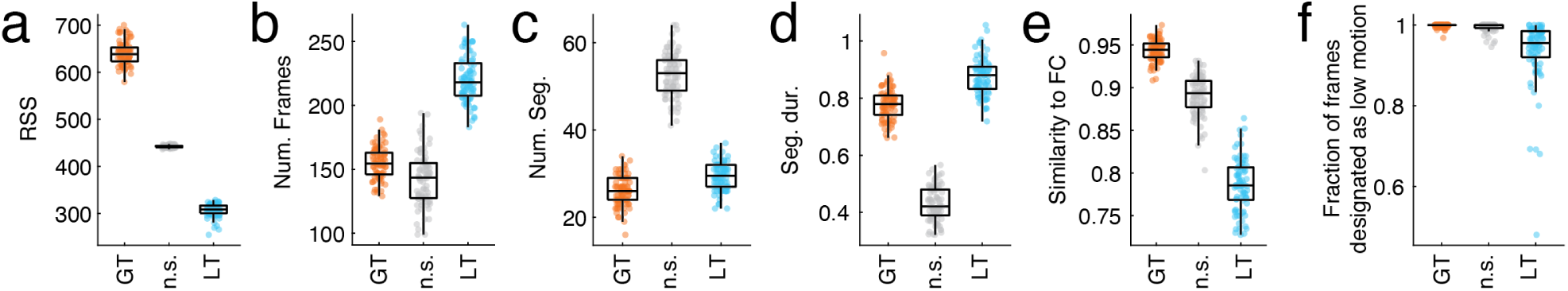
Statistical comparison of high- and low-amplitude frames using MyConnectome dataset. In the main text, we described a statistical procedure for partitioning frames into three categories based on how their RSS values compared to those of a null model. We applied that model to data from the MyConnectome Project and compared the three categories across multiple features, including their RSS amplitude, number of frames associated with each category, the number of contiguous frames of the same category (we call these “sequences”), sequence duration, the similarity of FC reconstructed using frames of each category to the static FC matrix, and the fraction of frames of a given category that were censored due to motion/data quality issues.

**FIG. S5.**
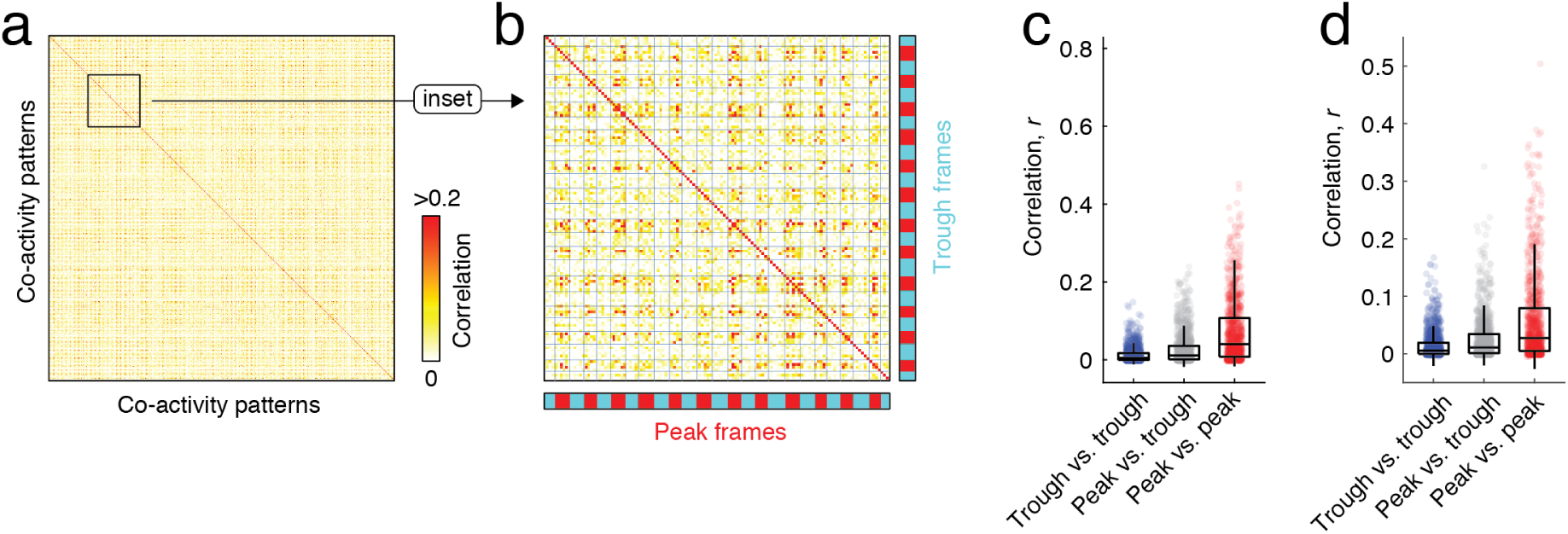
Cross-scan similarity of cofluctuation patterns in MyConnectome data and Midnight Scan Club data with individualized parcels. In the main text we demonstrated that peak cofluctuation patterns were repeated across scans, while trough cofluctuation were not. Here, we replicate this finding using data from 84 resting-state scans from the MyConnectome data. (*a*) Similarity matrix of cofluctuation patterns, including both peaks and troughs. (*b*) We show a smaller section of the complete similarity matrix in greater detail to highlight the relationship between peak and trough similarity. (*c*) Boxplot of similarity scores, aggregated based on whether the similarity was computed between two trough patterns, a trough and a peak pattern, or a peak and a peak pattern. We also show that we obtain a similar effect when using individualized parcellations of participant’s brains in the Midnight Scan Club. As in panel *c*, panel *d* depicts similarity scores, aggregated based on whether the similarity was computed between two trough patterns, a trough and a peak pattern, or a peak and a peak pattern.

**FIG. S6.**
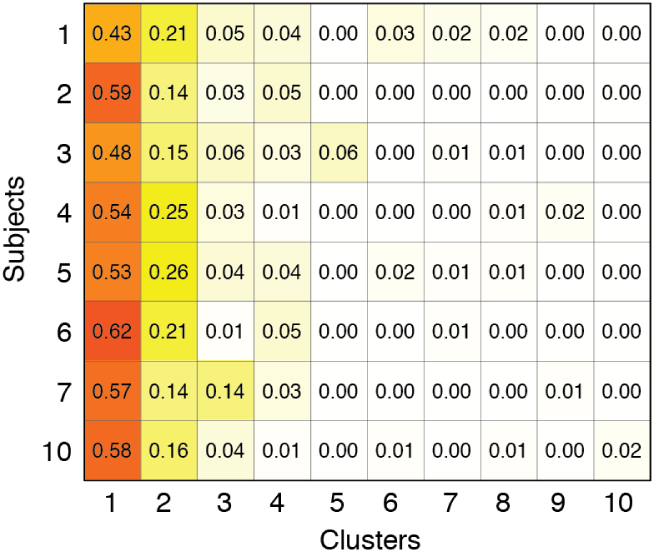
Community frequency by subject. Here, we show the frequency with which subjects (rows) visit the first ten communities (columns).

**FIG. S7.**
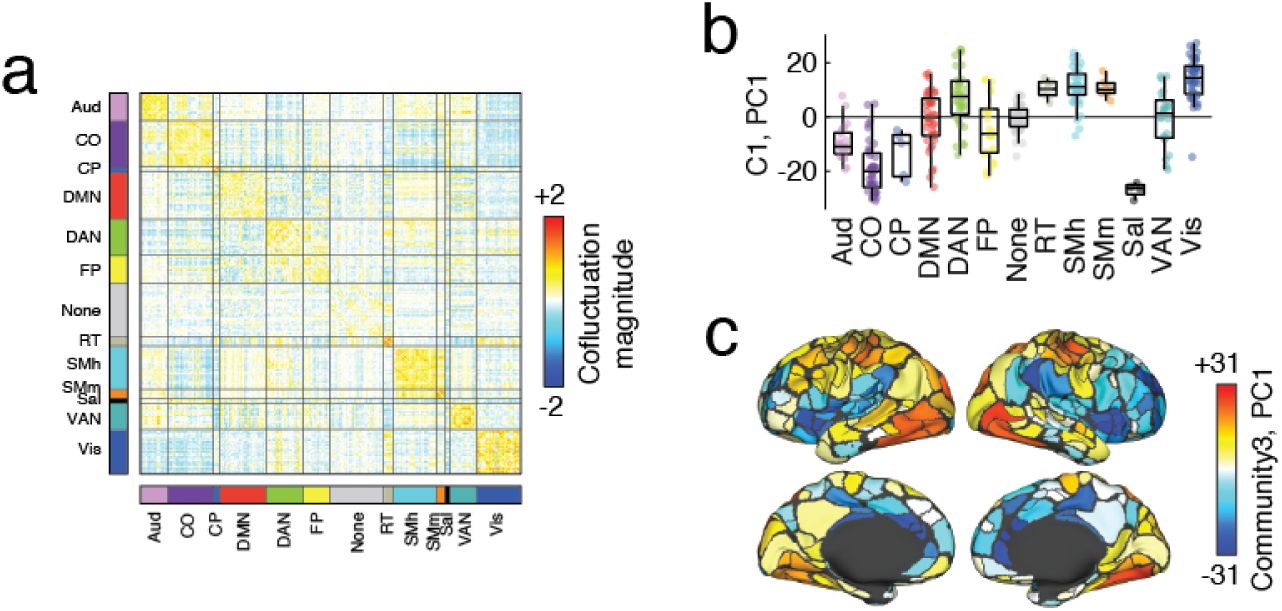
Summary of community 3 as an aggregated community. In the main text, we described the two large, cohesive communities that resulted from clustering peak cofluctuation patterns. The clustering algorithm also generated a large number of smaller communities. Only a fraction of participants were represented in each of these communities (there were no cases where all participants were represented). For simplicity, we grouped the smaller communities and treated them as if they had the same community label (community 3). Here, we summarize some features of that community. (*a*) cofluctuation pattern. We applied PCA to the fMRI BOLD time series associated with each peak cofluctuation pattern. In panel *b*, we show the first component divided into brain systems. (*c*) The same component projected onto the cortical surface.

**FIG. S8.**
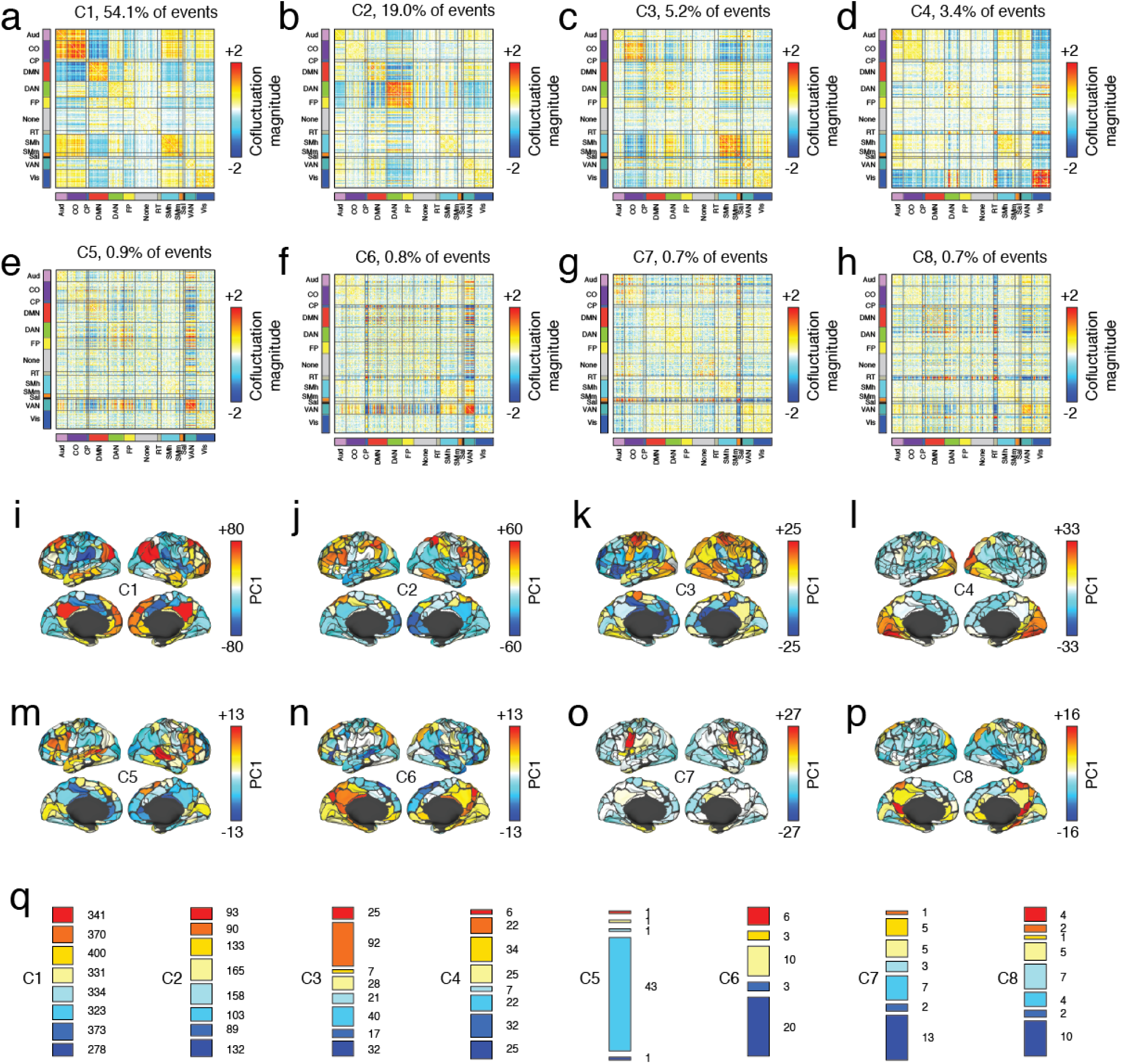
Summary of the individual communities that comprise community 3. In the main text, we described the two large, cohesive communities that resulted from clustering peak cofluctuation patterns. The clustering algorithm also generated a large number of smaller communities. In the main text we grouped these smaller communities into a single community labeled ‘3’. Here, we examine those smaller communities in greater detail along with communities 1 and 2, for completeness. In panels *a* -*h*, we show cofluctuation patterns for eight communities that included data from multiple participants and accounted for at least 0.5% of all detected events. Above each matrix, we show the fraction of all events accounted for by that community. In panels *i* -*p*, we show the activity patterns that underpin each community, plotting the first principal component of corresponding activity on the surface of the brain. In panel *q*, we show participant composition of each community. Here, different colors correspond to different participants. The height of each bar indicates the relative number of events assigned to a given community from a given participant.

**FIG. S9.**
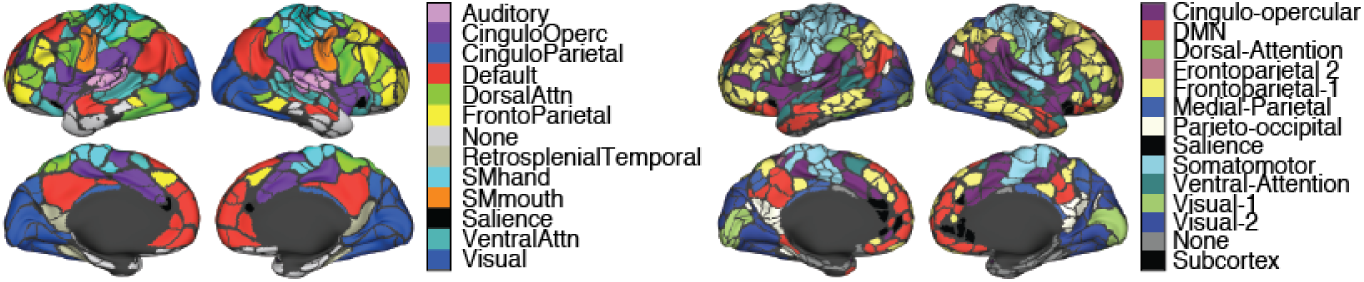
Brain systems projected onto cortical surface. (*left*) Gordon atlas. (*right*) MyConnectome atlas.

**FIG. S10.**
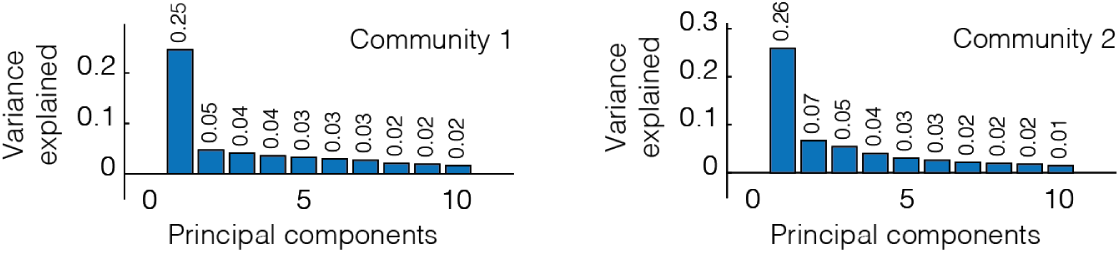
Variance explained by additional components. In the main text we applied PCA to activity patterns corresponding to peak cofluctuation. We reported variance explained by the first component for communities 1 and 2. Here, we show explained variance for the next nine components. Note that there is a sharp reduction in variance explained after the first component.

**FIG. S11.**
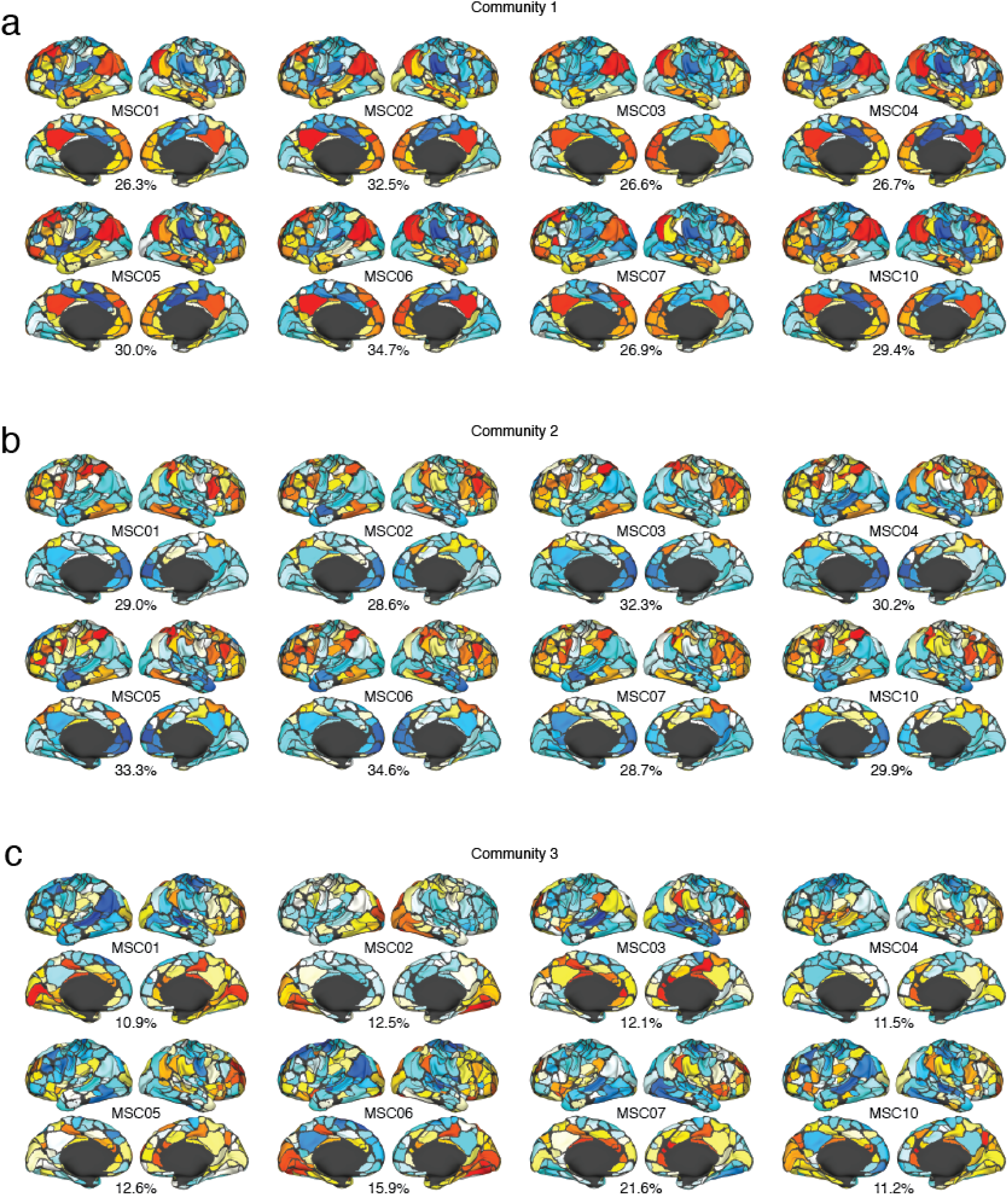
Participant-level PCA analysis of peak activity patterns. In the main text we, applied PCA to activity patterns corresponding to peak cofluctuation. Those analyses were performed at the group level. Here, we repeat this analysis at the level of individual participants. Panels *a*-*c* show PC1 for each of the three communities. The text below each surface plot indicates the percent variance explained by that PC.

**FIG. S12.**
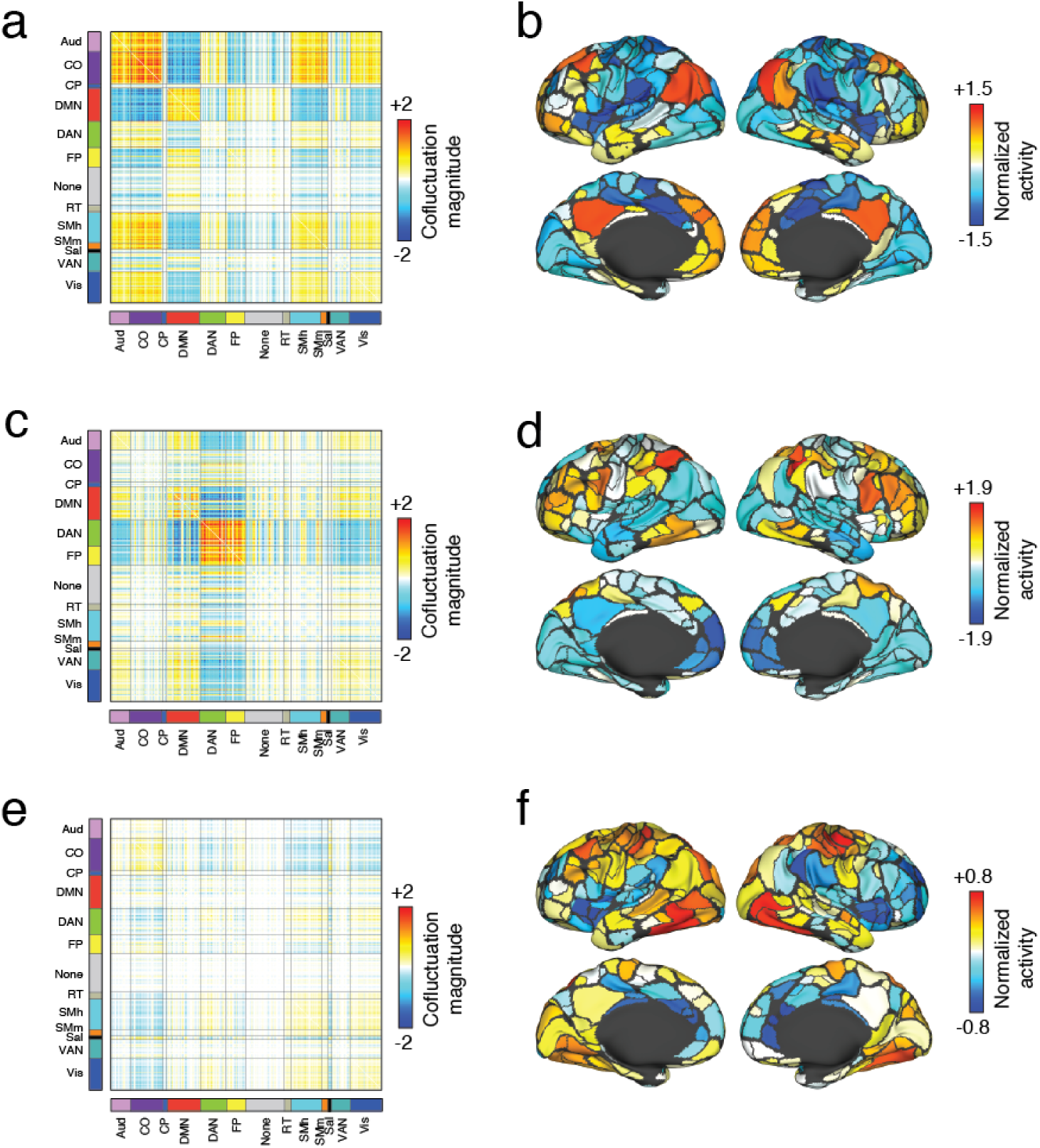
Alternative method for estimating cofluctuation and activity patterns. In the main text we identified three recurring patterns of cofluctuation and used a PCA-based method to identify their corresponding modes of activity. However, the cofluctuation between nodes *i* and *j* must satisfy for all {*i, j*} the following relationship: *C*_*ij*_ = *z*_*i*_*z*_*j*_, where *z*_*i*_ is the activity level of region *i*. In general, PCA will generate modes of activity that do not satisfy this relationship. To estimate this pattern, we used a greedy algorithm to identify the vector **z** = [*z*_1_, …, *z*_*N*_] whose cofluctuation matrix **C** = {*C*_*ij*_} minimizes the distance (root sum of squares) between the community-representative centroid. In panels *a, c*, and *e*, we show the cofluctuation matrices for communities 1, 2, and 3. Panels *b, d*, and *f* depict the activity pattern used to estimate cofluctuation.

**FIG. S13.**
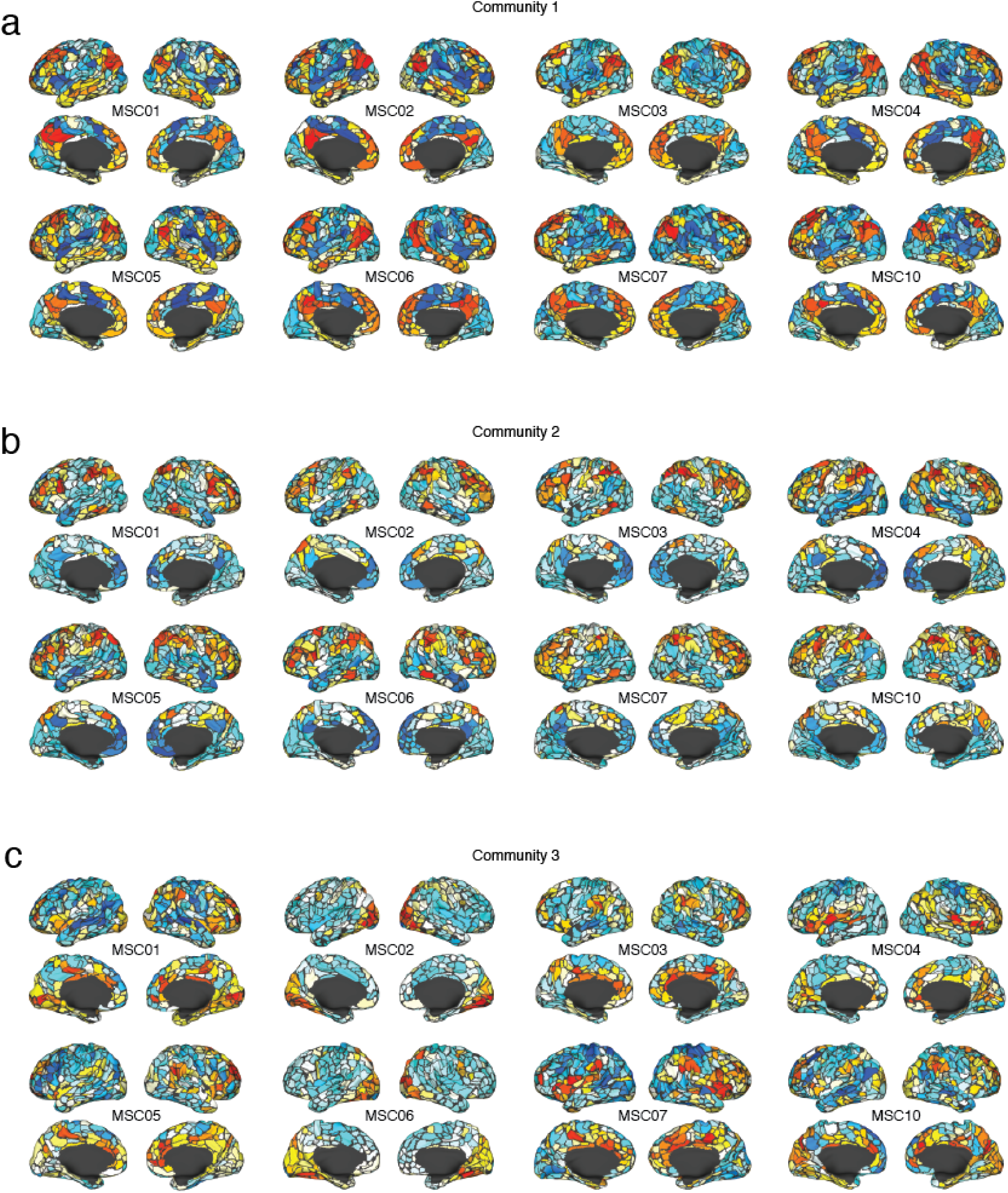
participant-level PCA analysis of peak activity patterns using individualized parcels. In the main text, we applied PCA to activity patterns corresponding to peak cofluctuation. Those analyses were performed at the group level using a common set of *N* = 333 brain regions. Here, we repeat this analysis at the level of individual participants using parcels fit individually to each participant. Panels *a*-*c* show PC1 for each of the three communities.

**FIG. S14.**
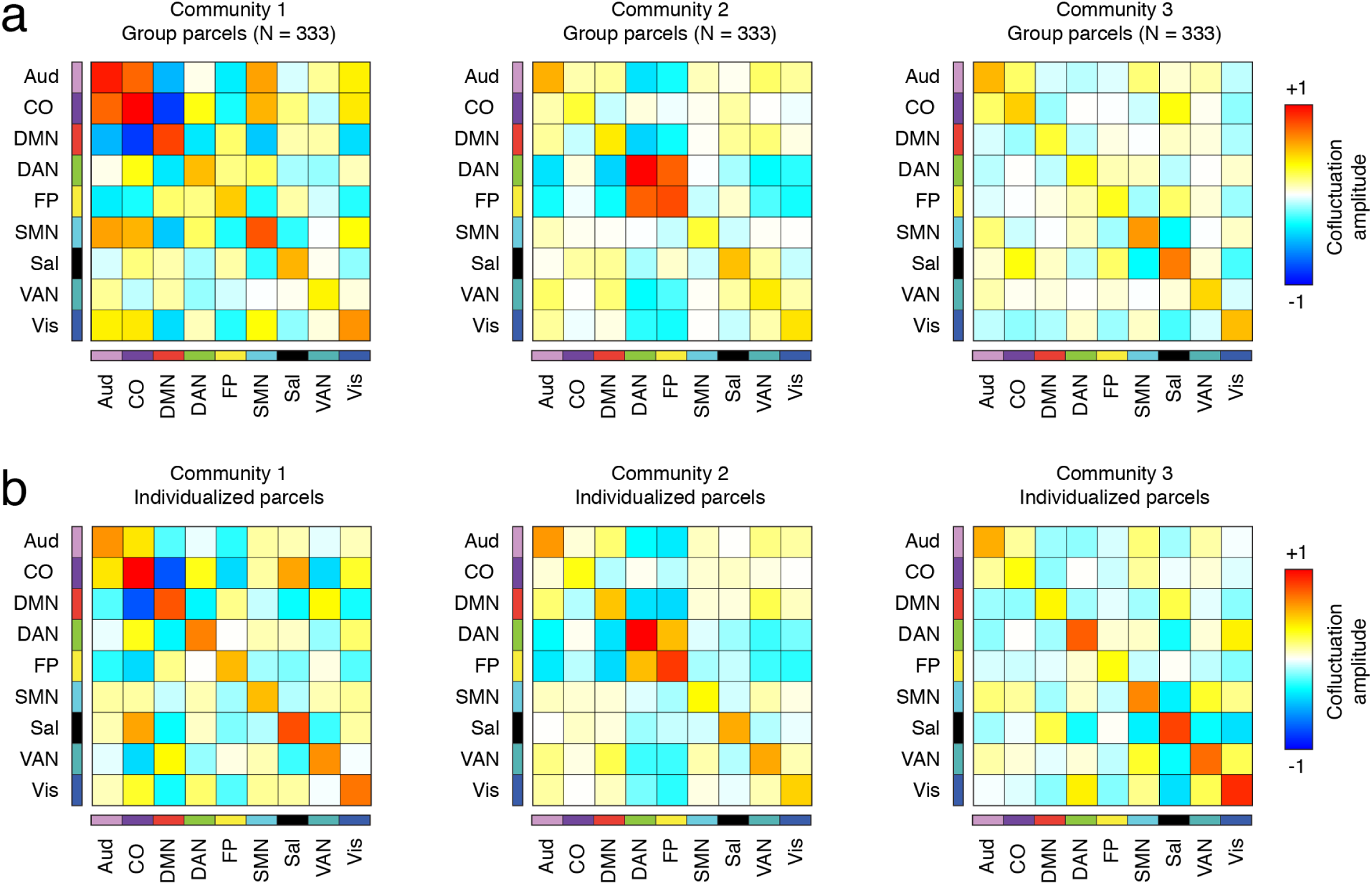
Similar system-level centroids using both group and individualized parcels. In the main text, we described three distinct cofluctuation patterns that were broadly shared across participants. Those patterns were estimated using a common set of *N* = 333 parcels. We also repeated those analyses using individualized parcels, which differed in number across participants. This variability made it impossible to compare cofluctuation patterns between participants as well as with the cofluctuation patterns estimated using the common set of parcels. To circumvent this issue, we estimated centroids at a system level by averaging the cofluctuation magnitude within and between nine systems that were expressed both at the group and individual level: Auditory, cingulo-opercular, default mode, dorsal attention, fronto-parietal, somatomotor, salience, ventral attention, and visual networks. Note, that with the group parcels the somatomotor network contains the somatmotor-hand and somatomotor-mouth networks. In the case of the individualized parcels, the somatomotor network contains the somatmotor-hand, somatomotor-foot, and somatomotor-mouth networks. Similarly, the visual network contains both lateral and primary visual networks for the individualized parcels. In general, we found a high level of correspondence between the two parcellations. The similarity values (Pearson correlation) of centroids for communities 1, 2, and 3 between datasets were *r* = 0.85, *r* = 0.97, and *r* = 0.84. In contrast, the similarity between centroids of *r* = 0.63±0.07. In *a* and *b* we show centroids estimated using the group and individualized parcels.

**FIG. S15.**
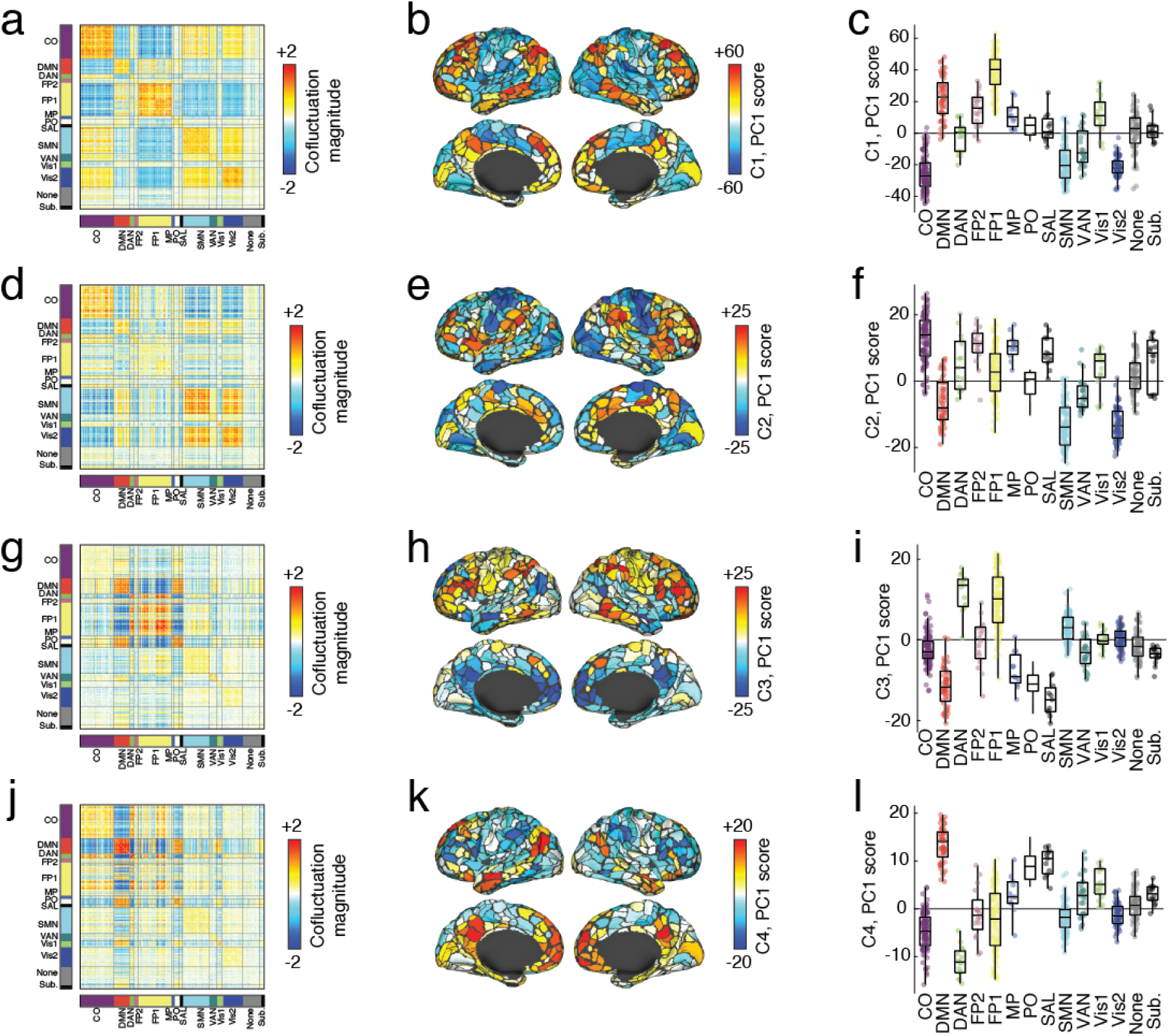
Results of multi-resolution consensus cluster applied to peak cofluctuation in MyConnectome dataset. In the main text we extracted peak cofluctuation patterns in the Midnight Scan Club dataset and applied a clustering algorithm to group patterns into communities. Here, we apply an identical analysis to MyConnectome data. The algorithm identified four large communities, which we summarize here. In panels *a, d, g*, and *j*, we show the mean cofluctuation for each community. Note that community 1 and community 4 in the MyConnectome data is nearly identical to community 1 in the Midnight Scan Club. Similarly, communities 2 and 3 bear strong resemblance to community 2 in the Midnight Scan Club analysis. Panels *b, e, h*, and *k* show the first principal component (PC1) of activity during each peak. Panels *c, f, i*, and *l* show boxplots of PCs sorted by system.

**FIG. S16.**
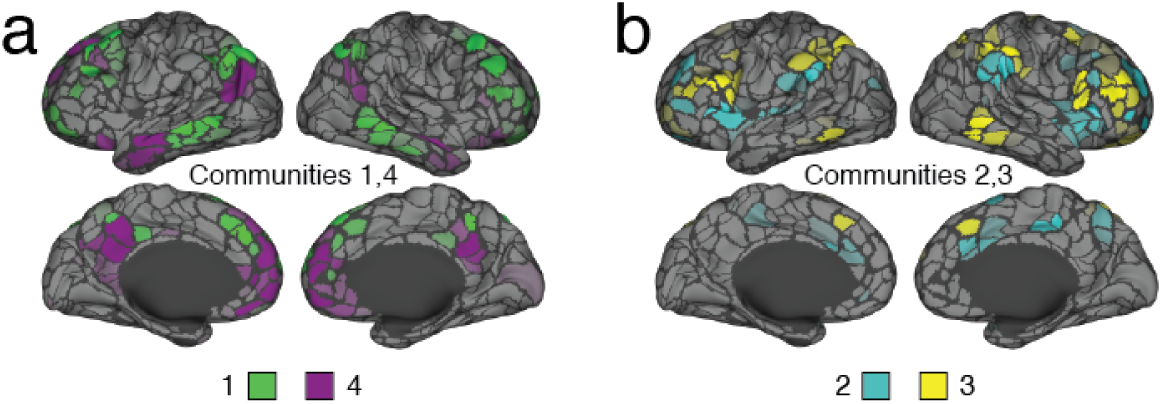
Interdigitated activity during peaks. We clustered cofluctuation during peaks and found that the four main communities we reported could be grouped into pairs that involved similar brain systems. Here, we show these pairs for interdigitated sub-divisions of canonical systems To do this, we calculated PC1 for each of the four communities. We then normalized PC1, dividing every element by 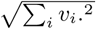, where *v*_*i*_ is the *i*th element (corresponding to region *i*) of PC1. This normalization ensures that the eigenvector has magnitude equal to unity. We then paired communities 1 with 4 and 2 with 3 and, for each region, we calculated which of the two normalized eigenvectors had a larger value. Here, we visualize the winners of the top quartile (25%) of regions. In *a* we show communities 1 and 4 and in *b* we show communities 2 and 3.

**FIG. S17.**
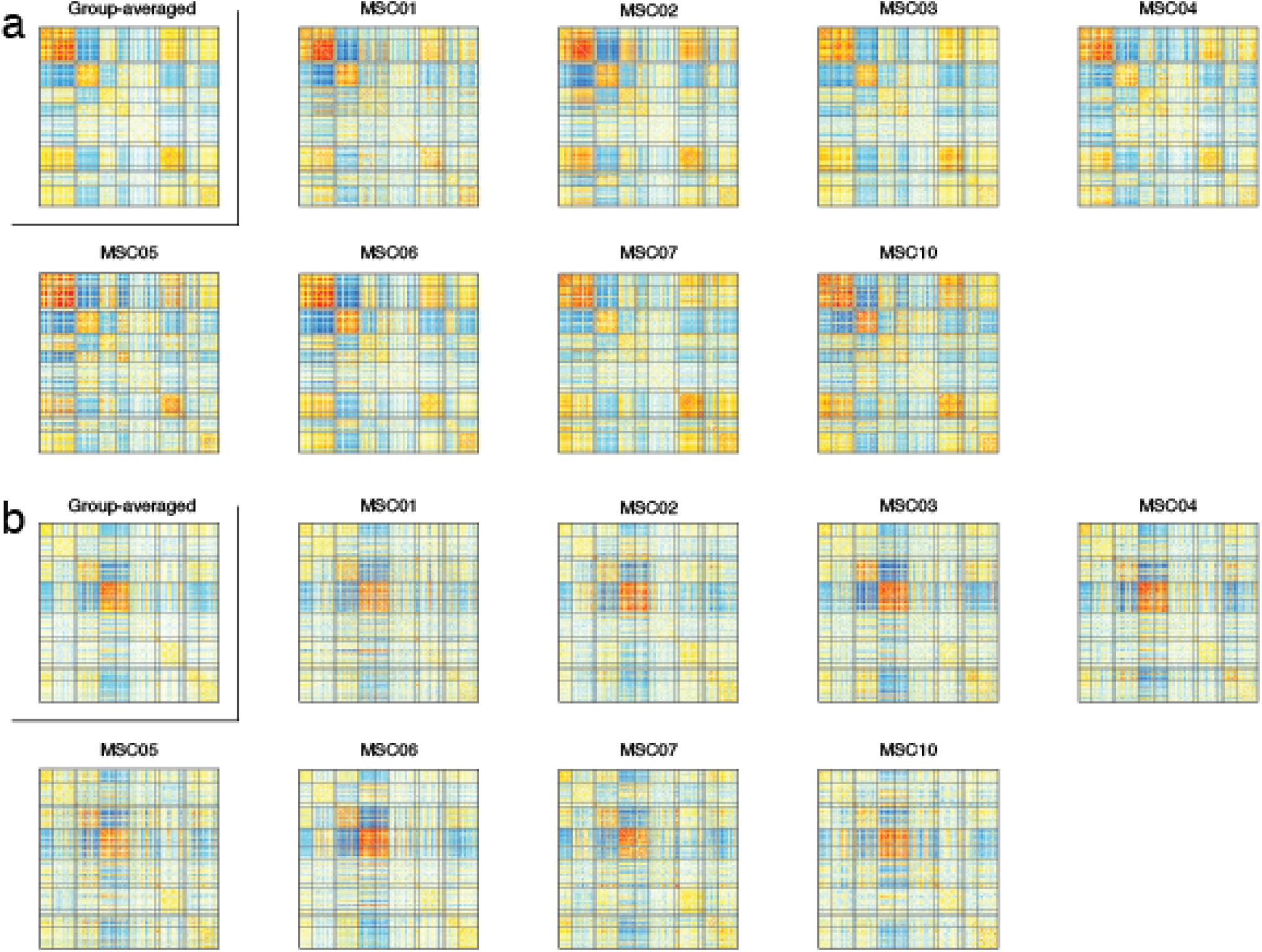
Group- and subject-level centroids. Matrix representations of group- and subject-level centroids for (*a*) community 1 and (*b*) community 2.

**FIG. S18.**
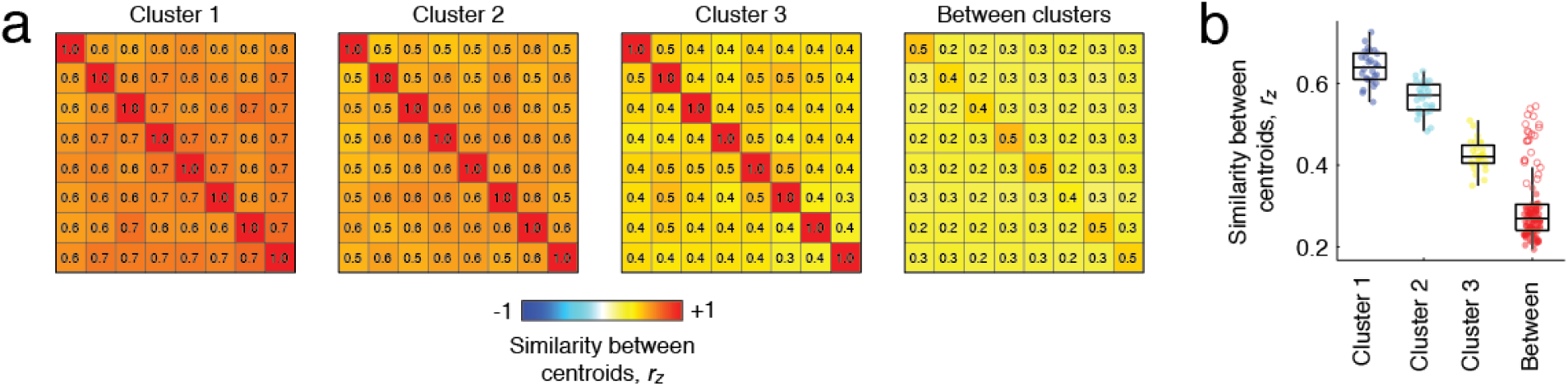
Within and between community similarity of centroids. In the main text we clustered cofluctuation peaks into three communities. Here, we calculated each participant’s centroid (mean cofluctuation pattern for each community) using only those peaks assigned to that community. (*a*) The similarity (Pearson correlation) of participants’ centroids to one another for communities 1, 2, and 3. We also show the similarity of centroids from different communities (*right*). (*b*) Similarity scores grouped by community. Note that in the “between” category, some points are solid and others outlined. The outlined points represent similarity scores from the same participant across different centroids.

**FIG. S19.**
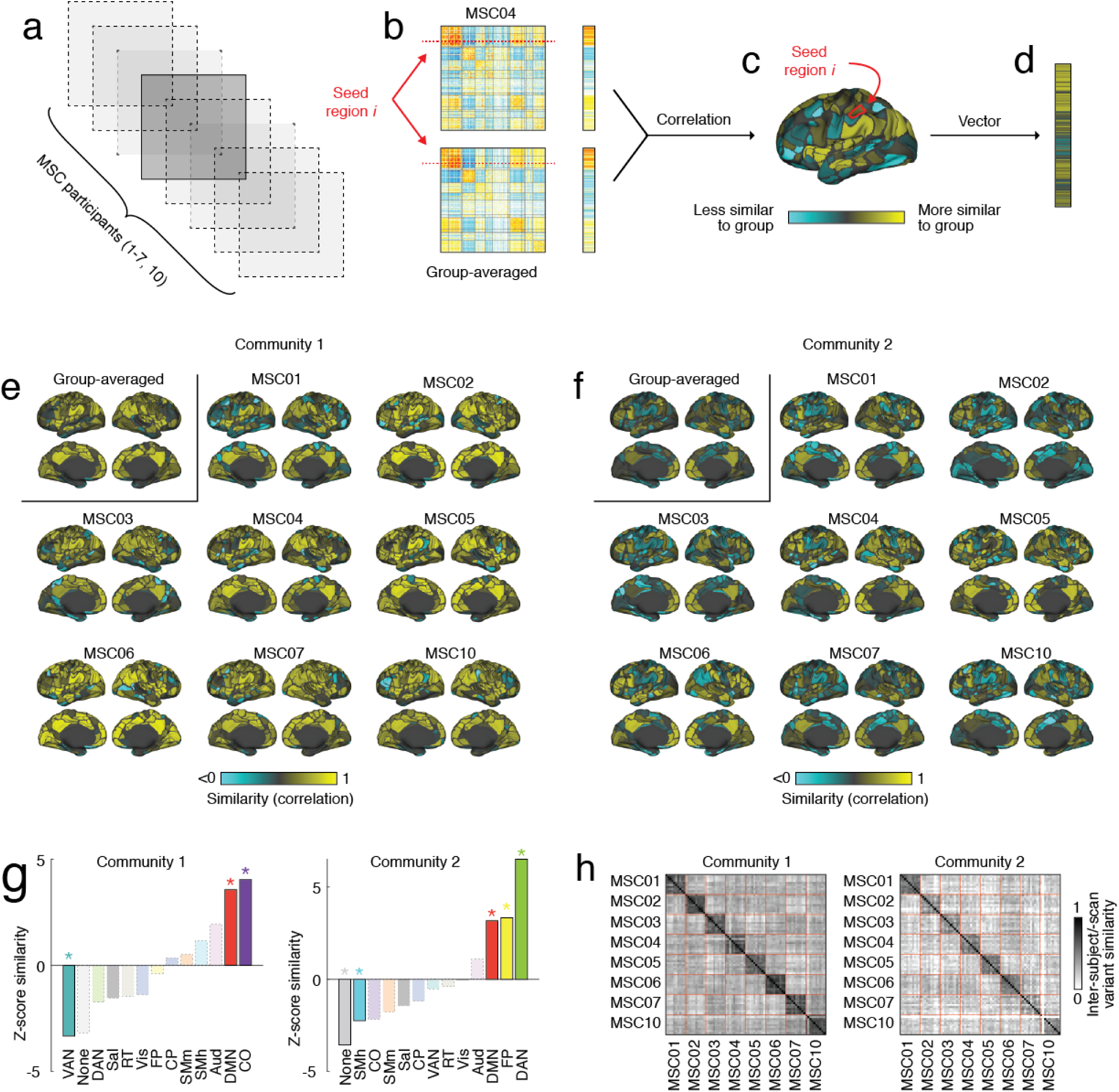
Identification and stability of subject-level deviations from group co-fluctuation patterns. In the main text we detected high-amplitude cofluctuations (events) and clustered them into two communities. Although later we used subject-level estimates of those community centroids in the predictive model, we characterized communities at the group level. Here, we describe subject-level deviations from the group-averaged centroids. (*a*) For a given community, we estimated community centroids for each subject and the group as a whole. (*b*) Separately for each subject and region, we calculated the similarity of its cofluctuation pattern with that of the group. (*c*) Repeating this procedure for each region yielded a whole-brain similarity map that we could project onto the cortical surface. (*d*) The map could also be vectorized and studied further. Panels *e* and *f* display group- and subject-level similarity maps for communities 1 and 2, respectively. For community level maps, we asked whether similarity values were clustered within specific brain systems. (*g*) We found that for community 1, the ventral attention network was, on average, more dissimilar than chance while the default mode and cingulo-opercular networks were more similar to the group than chance. For community 2, the regions lacking a clear system assignment (‘none’) and the somatomotor-hand system were more dissimilar than expected, while default mode, fronto-parietal, and dorsal attention networks were more similar. (*h*) We assessed the extent to which the subject-level similarity maps were stable within and between individuals by computing the pairwise similarity matrix of maps across all scans and subjects. In general, we found higher levels of similarity within subjects than between.

**FIG. S20.**
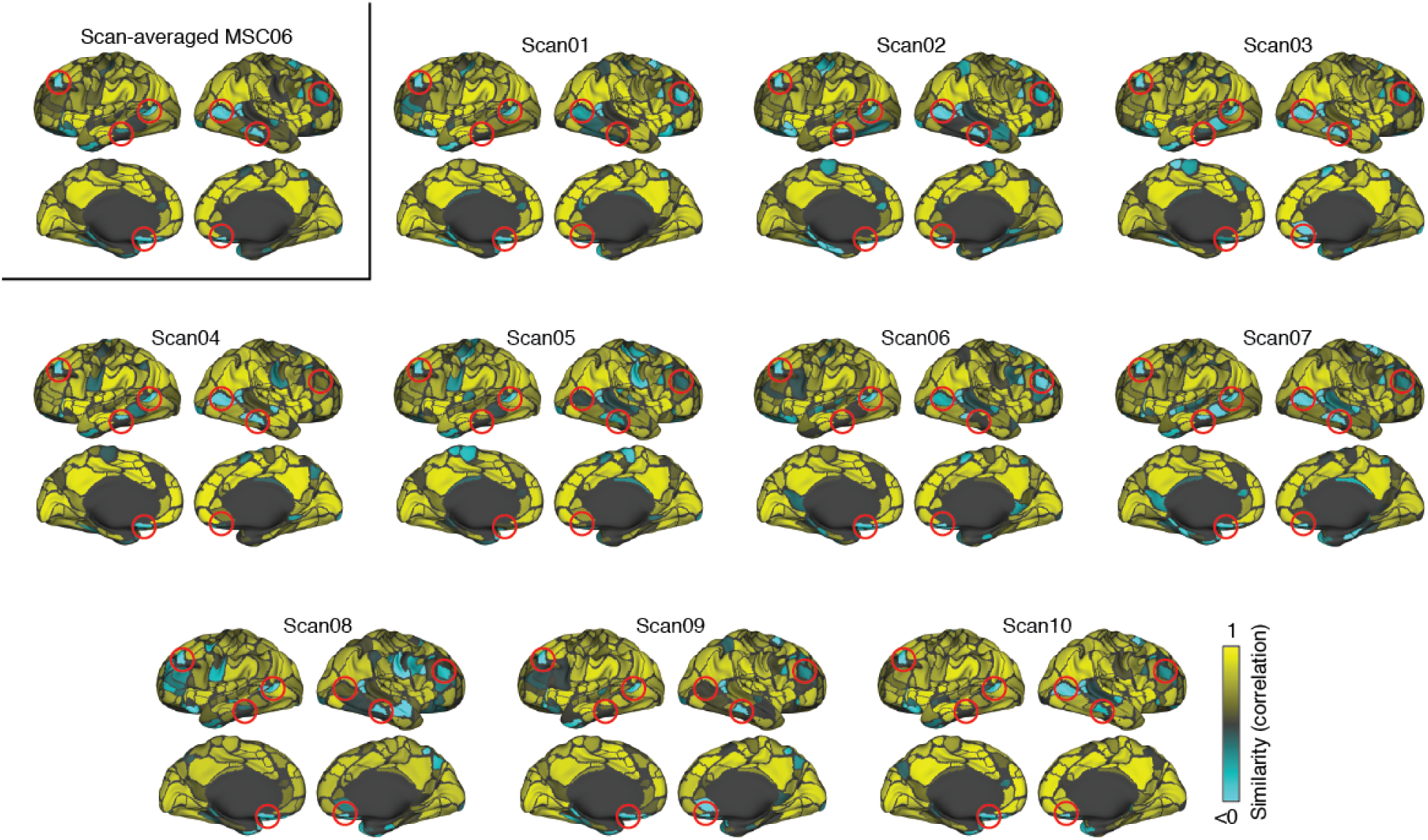
Example similarity maps for MSC06. Here, we show the regional similarity maps of data from MSC06 the group-averaged community 1 centroid. In each plot, we highlight patterns of dissimilarity (blue regions) that are consistently expressed across scan sessions.

**FIG. S21.**
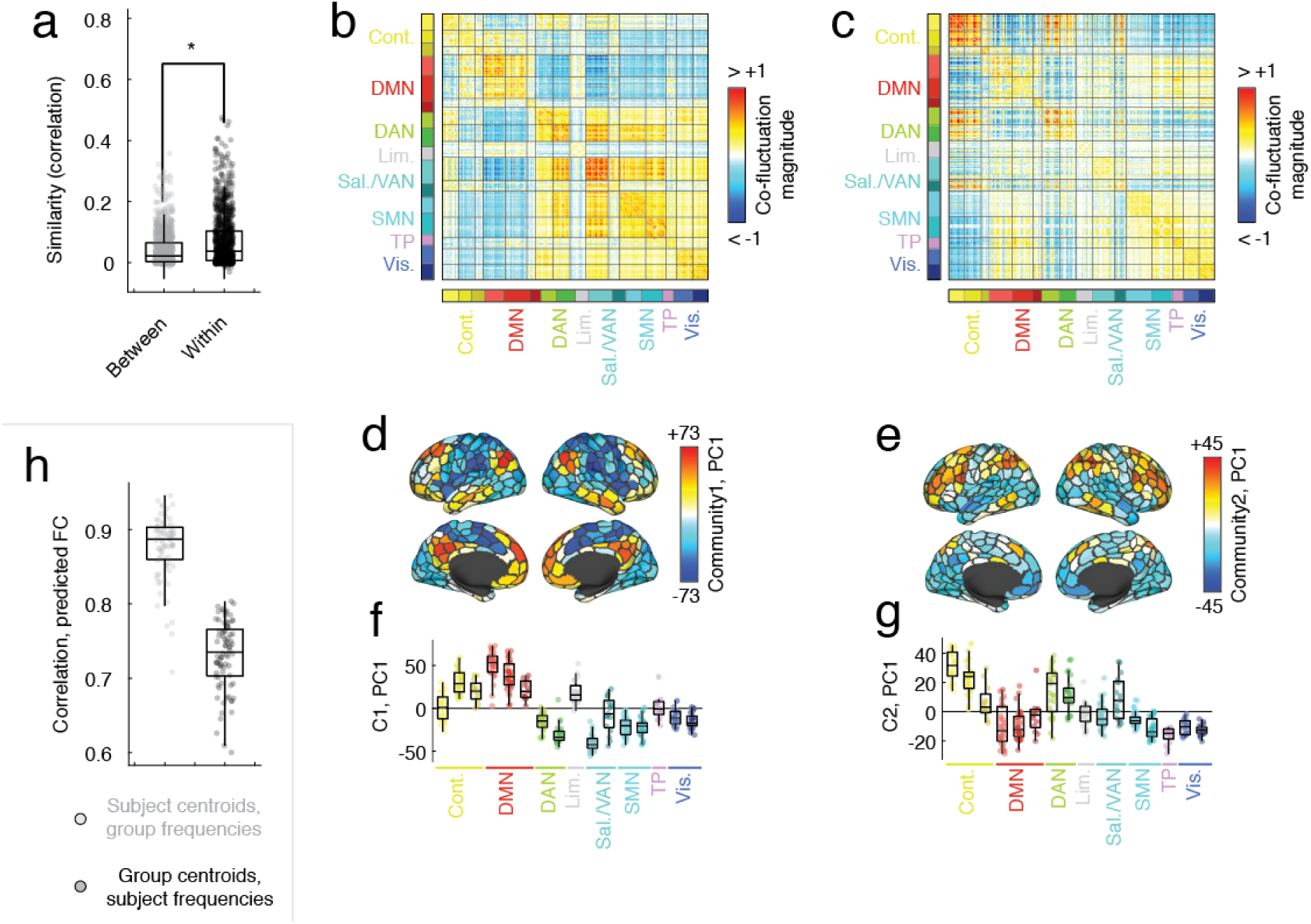
Replication of main results using alternative processing pipeline. In the main text, we show that events detected using MSC data are more similar within-subjects than between, can be clustered into two communities, and that subject-level variants of these communities along with group-level frequencies lead to accurate predictions of FC. Here, we replicate these findings after reprocessing the MSC data using an alternative pipeline [42] and parcellating the brain into *N* = 400 cortical regions [43]. (*a*) Similarity of high-amplitude co-fluctuations within and between subjects. Within-subject similarity is significantly greater than between-subject similarity (*t*-test; *p <* 10^−15^). Panels *b* and *c* show group-level estimates of community centroids. Note that here the event and community detection analyses were redone using the newly processed data. Panels *d* and *e* depict the first principal component of event activity projected onto cortical surface while panels *f* and *g* show the same data as a boxplot. (*h*) Results of predictive model comparing H1 and H2. As in the main text, the model with subject-specific centroids outperformed the model with subject-specific frequencies (paired sample *t*-test, *p <* 10^−15^).

**FIG. S22.**
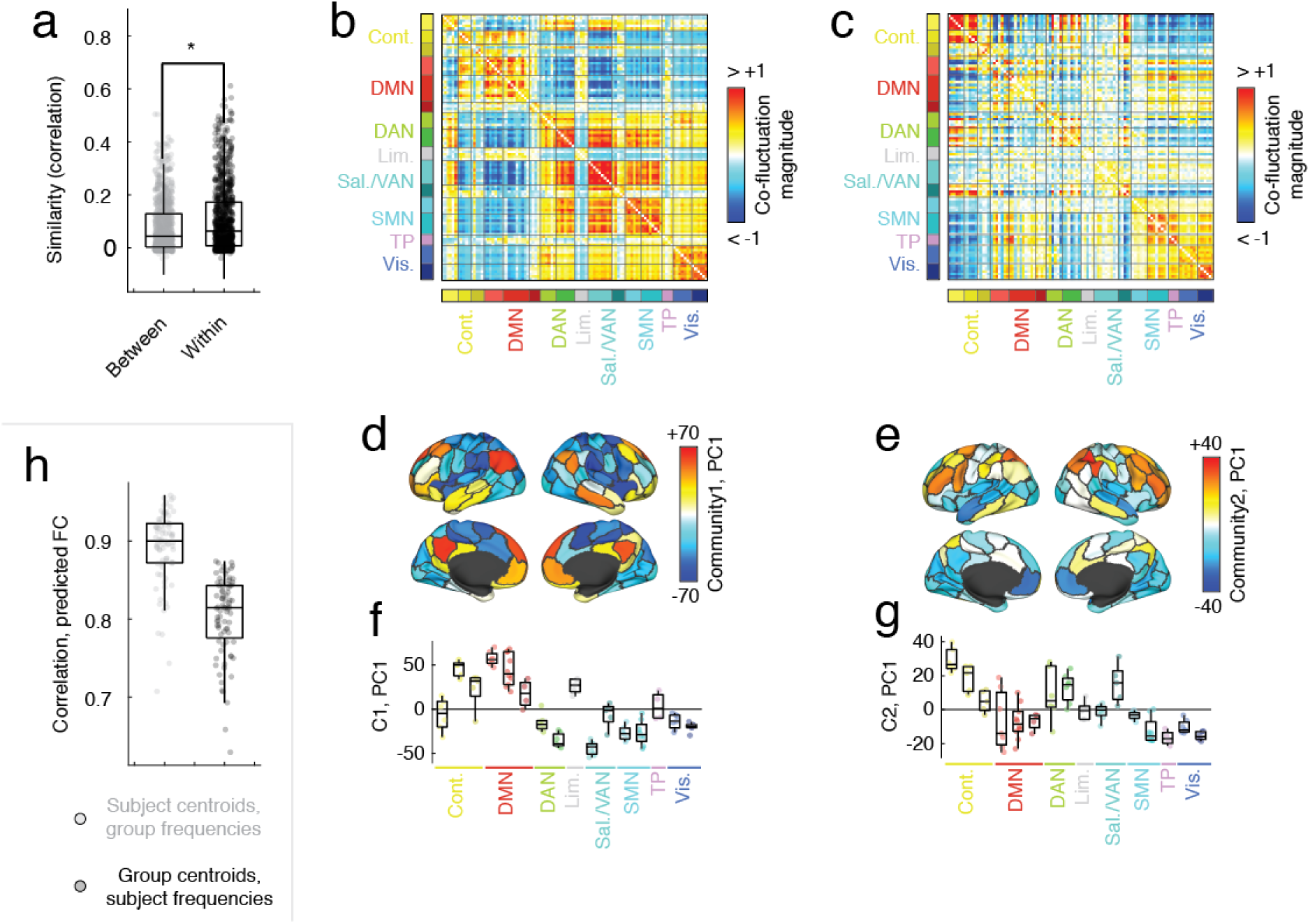
Replication of main results using coarser cortical parcellation. In the main text, we show that events detected using MSC data are more similar within-subjects than between, can be clustered into two communities, and that subject-level variants of these communities along with group-level frequencies lead to accurate predictions of FC. In the previous figure, we replicate these findings after reprocessing the MSC data using an alternative pipeline [42] and parcellating the brain into *N* = 400 cortical regions [43]. Here, we further replicate those findings using a coarser parcellation of the brain into *N* = 100 regions. (*a*) Similarity of high-amplitude co-fluctuations within and between subjects. Within-subject similarity is significantly greater than between-subject similarity (*t*-test; *p <* 10^−15^). Panels *b* and *c* show group-level estimates of community centroids. Note that here the event and community detection analyses were redone using the newly processed data. Panels *d* and *e* depict the first principal component of event activity projected onto cortical surface while panels *f* and *g* show the same data as a boxplot. (*h*) Results of predictive model comparing H1 and H2. As in the main text, the model with subject-specific centroids outperformed the model with subject-specific frequencies (paired sample *t*-test, *p <* 10^−15^).

**FIG. S23.**
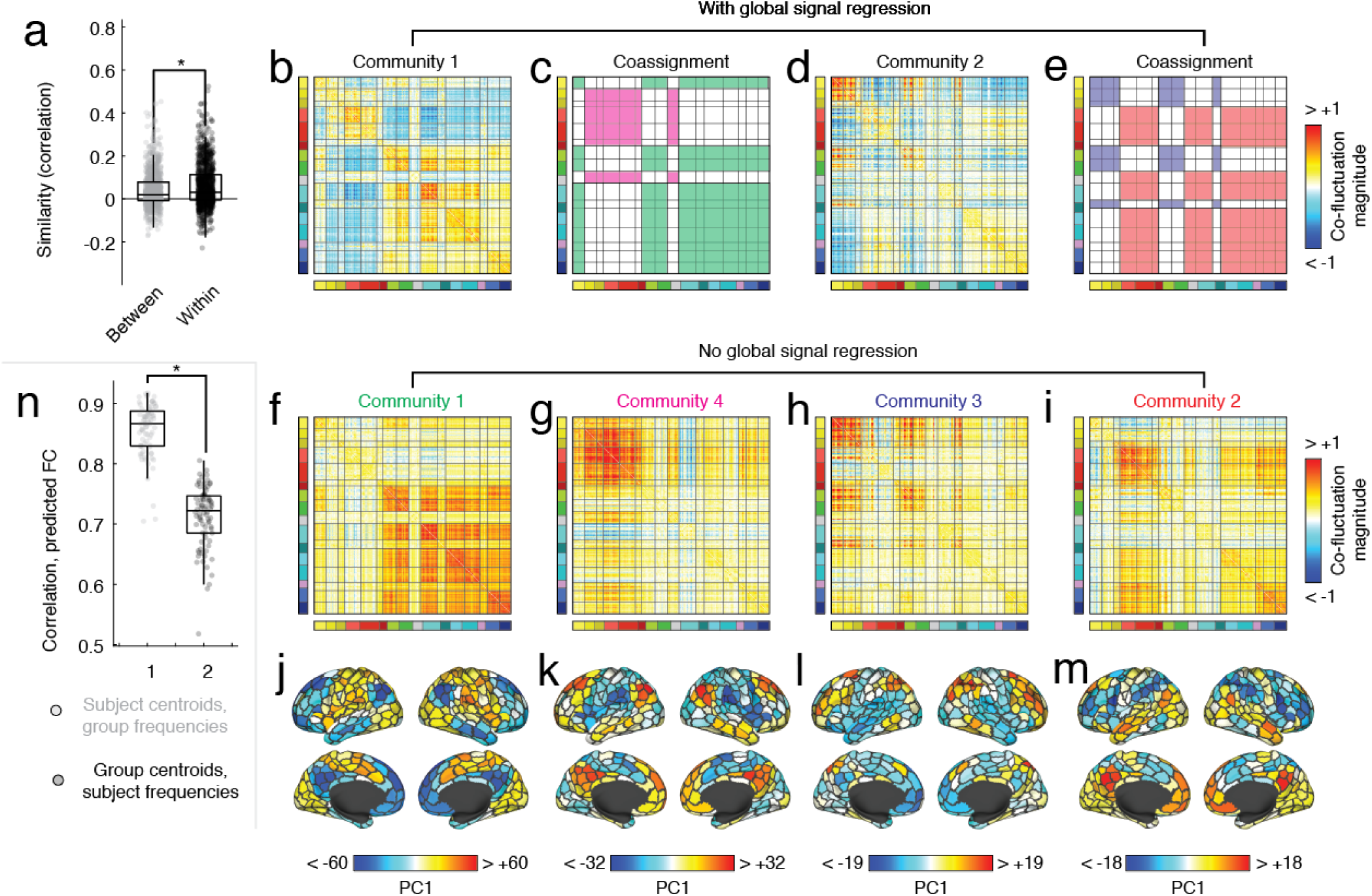
Replication of main results using alternative processing pipeline with no global signal regression. In the main text, we show that events detected using MSC data are more similar within-subjects than between, can be clustered into two communities, and that subject-level variants of these communities along with group-level frequencies lead to accurate predictions of FC. In the previous two figures, we replicate these findings after reprocessing the MSC data using an alternative pipeline [42] and parcellating the brain into *N* = 100 and *N* = 400 cortical regions [43]. Here, we further replicate those findings after excluding global signal regression from the processing pipeline by using the anatomical CompCor method [100]. (*a*) Similarity of high-amplitude co-fluctuations within and between subjects. Within-subject similarity is significantly greater than between-subject similarity (*t*-test; *p <* 10^−15^). Without global signal regression, the previously described anticorrelations in high-amplitude co-fluctuation patterns are not as evident. In panels *b* and *d* we show those communities for reference, highlighting the anticorrelated groups of brain regions in *c* and *e* and designating the groups with distinct colors. Without global signal regression, we find that each of these anticorrelated groups of regions forms its own community. In panels *f* -*i* we show the centroids of these communities. The labels above each matrix are colored according to which of the anticorrelated groups they most closely resemble. Note that these four communities were also the four largest. Panels *j* -*m* project onto the cortical surface the first principal component of the underlying brain activity for each of the four communities. (*n*) Results of predictive model comparing H1 and H2. As in the main text, the model with subject-specific centroids outperformed the model with subject-specific frequencies (paired sample *t*-test, *p <* 10^−15^).

**FIG. S24.**
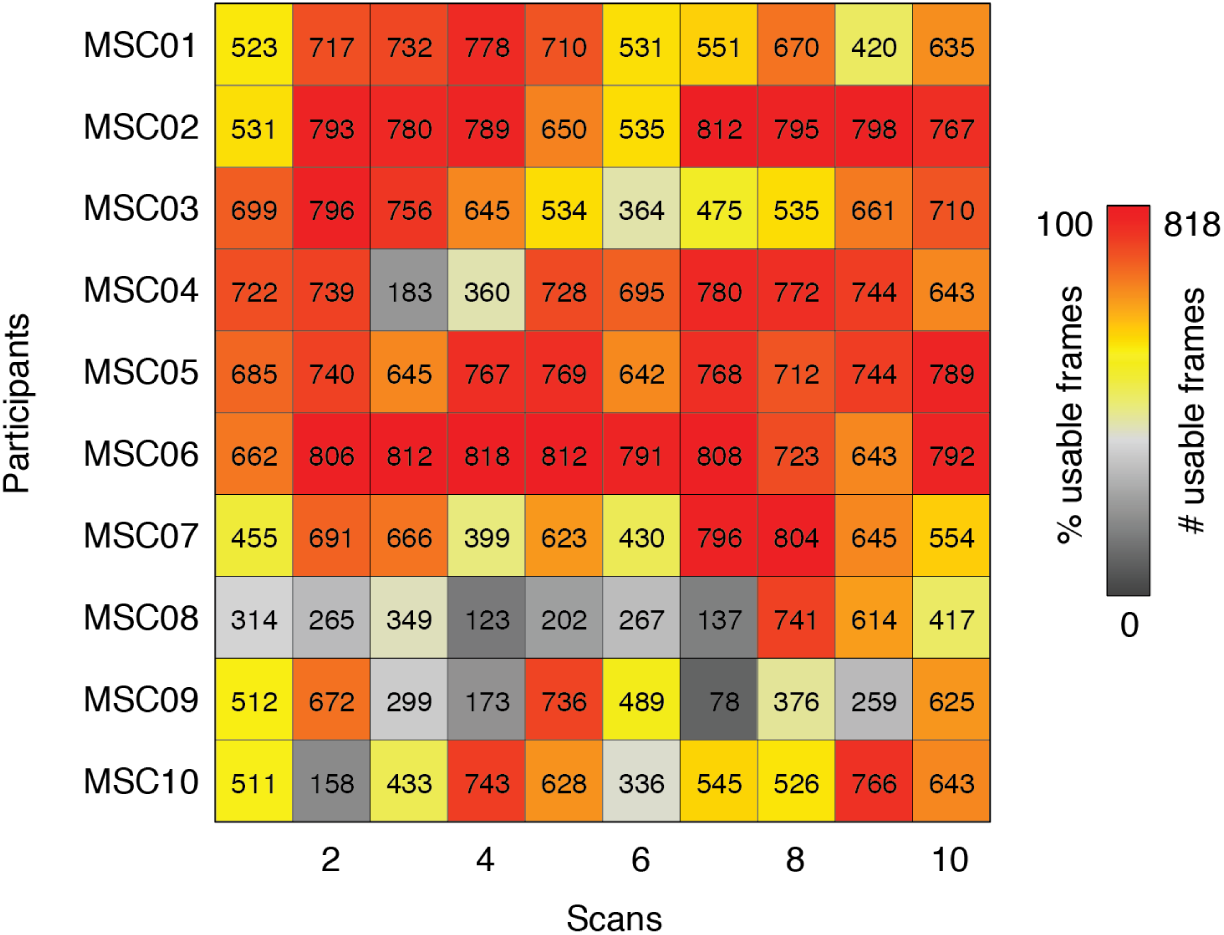
Summary of usable data by subject and scan. Here, we report the number of usable frames for each participant and scan. Note that we exclude MSC08 and MSC09 from analyses given that in more than half of their scans, more than 50% of the frames were discarded.

